# The structure of simple satellite variation in the human genome and its correlation with centromere ancestry

**DOI:** 10.1101/2023.07.03.547555

**Authors:** Iskander Said, Daniel A. Barbash, Andrew G. Clark

## Abstract

Although repetitive DNA forms much of the human genome, its study is challenging due to limitations in assembly and alignment of repetitive short-reads. We have deployed *k-Seek*, software that detects tandem repeats embedded in single reads, on 2,504 human genomes from the 1,000 Genomes Project to quantify the variation and abundance of simple satellites (repeat units < 20 bp). We find that homopolymers and the *Human Satellite 3* monomer make up the largest portions of simple satellite content in humans (mean of ∼19 Mb combined). We discovered∼50,000 rare tandem repeats that are not detected in the *T2T-CHM13v2.0* assembly, including undescribed variants of telomeric- and centromeric repeats. We find broad homogeneity of the most abundant repeats across populations, except for AG-rich repeats that are more abundant in African individuals. We also find cliques of highly similar AG- and AT-rich satellites that are interspersed and form higher-order structures that covary in copy number across individuals, likely through concerted amplification via unequal exchange. Finally, we use centromere-linked polymorphisms to estimate centromeric genetic relatedness between individuals and find a strong predictive relationship between centromeric lineages and centromeric simple satellite abundances. In particular, *Human Satellite 2* and *Human Satellite 3* abundances correlate with clusters of centromeric ancestry on chromosome 16 and chromosome 9, with some clusters structured by population. These results provide new descriptions of the population dynamics that underlie the evolution of simple satellites in humans.

## Introduction

Repetitive DNA is a near-ubiquitous feature of eukaryotic genomes and in humans represents ∼54% of the genome (Hoyt et al. 2022). Repetitive DNA in the human genome is ∼89% interspersed transposable element sequences and ∼11% satellites, tandemly repeating arrays of homogenized sequence motifs with individual arrays ranging in length from a few hundred base pairs to several megabases (Altemose et al. 2022; Hoyt et al. 2022). These tandem repeats are diverse in sequence, implicated in human disease etiology, and some form essential genomic components, yet the extent of their variation in sequence and abundance in human populations is poorly understood (Mirkin 2007; Brand and Levine 2021; Altemose et al. 2022). Tandem repeats can be divided into three classes. Microsatellites (also known as STRs and VNTRs) are made up of short sequence motifs, or monomers, (typically, 1-6 bp) that form arrays of a few hundred base pairs that are interspersed throughout the genome (Ellegren 2004). Microsatellites can be in the coding regions of essential genes or regulatory regions and their copy-number changes are implicated in several diseases and disorders, such as Huntington’s disease, gout and autism spectrum disorder (Pearson et al. 2005; Mirkin 2007; Mitra et al. 2021; Mukamel et al. 2021).

The other classes of tandem repeats are simple and complex satellites, which we define as being made up of short (<20 bp) or long (>20 bp) monomers, respectively, repeated in long tandem arrays that can span up to several Mb. Despite the abundance of satellites in the genome, it remains unknown what proportion have any biological function. Some satellites can form essential structural components of the telomeres and centromeres of human and other eukaryotic chromosomes (Brand and Levine 2021). Human centromeres are composed of a mixture of long tandem arrays of simple satellites (*e.g. Human Satellite 2,3*) and complex satellites (*e.g. Alpha Satellites*), while telomeres are composed primarily of long arrays of a hexameric simple satellite (Prosser et al. 1986; Allshire et al. 1989; Lee et al. 1997; Altemose et al. 2022). These satellites, particularly those of the centromere, can form higher-order structures where variant monomers are interspersed with each other in specific periodicities to create substructures of multiple repeat families and subfamilies within larger arrays (Willard and Waye 1987; Durfy and Willard 1989; Altemose et al. 2014; Altemose et al. 2022).

Some simple satellites are transcribed into long non-coding RNAs that have roles in spermatogenesis, chromatin regulation, heat shock response, and cell division (Goenka et al. 2016; Mills et al. 2019; Yadav et al. 2020; Landers et al. 2021; Novo et al. 2022). However, many satellite repeats may not have specific regulatory functions, but rather affect gene expression in aggregate through the sequestration of heterochromatic proteins and alteration of chromatin states in *trans* (Dimitri and Pisano 1989; Zhou et al. 2012; Berloco et al. 2014; Francisco and Lemos 2014; Kelsey and Clark 2017; Delanoue et al. 2023). Variation of satellites may also impact their effects. For example, sequence variants of *Alpha Satellite* repeats have differential ability to nucleate the formation of the kinetochore, and differential positioning of the kinetochore on the X chromosome has been observed within human populations (Aldrup-MacDonald et al. 2016; Altemose et al. 2022). Additionally, the variation of both the amount and sequence of constitutive heterochromatin in the Y-chromosome affects gene expression and chromatin state (Zhou et al. 2012; Berloco et al. 2014; Kelsey and Clark 2017; Delanoue et al. 2023). However, the full extent of satellite variation in both abundance and sequence and how much of this variation affects organismal fitness have yet to be fully explored.

Many satellites are rapidly evolving. For example, *Alpha Satellite* has conserved biological function across taxa, yet shows rapid rates of evolution in sequence and abundance (Henikoff et al. 2001). There are highly divergent species-specific simple satellites and satellite subfamilies in closely related species of *Drosophila* and primates that have rapidly evolved in sequence and abundance (Waye and Willard 1989; Haaf and Willard 1997; Jarmuz et al. 2007; Wei et al. 2018; Cechova et al. 2019). Species-specific satellites are even implicated in hybrid incompatibility in *Drosophila* species (Bayes and Malik 2009; Ferree and Barbash 2009; Satyaki et al. 2014). There is also considerable intraspecific variation of simple satellite abundances in natural and experimental populations of *D. melanogaster*, *Chlamydomonas reinhardtii* and *Daphnia pulex* (Wei et al. 2014; Flynn et al. 2017; Flynn et al. 2018). These observations are somewhat contradictory to the theoretical models of concerted satellite evolution, which predict that within a given species satellite sequences and variation should be homogenized (Smith 1976; Perelson and Bell 1977; Stephan 1989; Stephan and Cho 1994). This deviation of the empirical data from the evolutionary models suggests that the evolutionary dynamics of simple satellites in populations are not well described and significant advancements to the population genetic models must be made to fully capture their evolutionary rates and variation, particularly if we aim to understand the selective forces acting on satellites.

Human population-genomics data provide a vast resource of thousands of high-quality genomes necessary to advance the evolutionary models of satellites. However, studies of the population variation of satellites in humans have been focused primarily on microsatellites (Payseur et al. 2011; Willems et al. 2014). The variation of simple and complex satellites has been studied to a lesser degree, but there is evidence of over 10-fold differences in *Human Satellite 3* abundance between human Y-haplogroups, 5-10 fold differences in centromere size, and tremendous diversity in the higher-order structures of centromeric repeats, pointing to rapid evolution of satellites within human populations (Altemose et al. 2014; Miga 2019; Suzuki et al. 2020; Altemose et al. 2022). Some satellite variation within the centromeres affects the formation and the positioning of the kinetochore, which may affect cell division, but it is unclear how much observed satellite variation affects function or organismal fitness (Aldrup-MacDonald et al. 2016; Altemose et al. 2022).

Studies of simple satellites in humans have been limited due to the technical challenges of aligning repetitive short-reads from Next Generation Sequencing (NGS) libraries. Recently there have been tremendous advances in assembling PacBio and Oxford Nanopore long-reads, allowing the Telomere-to-Telomere (T2T) consortium to fully assemble the human genome, including the telomeres and centromeres that were notably absent in previous assemblies (Suzuki et al. 2020; Ebert et al. 2021; Nurk et al. 2022; Porubsky et al. 2022). Studies on the *T2T-CHM13v2.0* genome assembly revealed patterns of layered expansions of repeats in the centromeres as well as variation in centromeric satellites within the X chromosomes of several individuals (Altemose et al. 2022). Additionally, pan-genome alignments of highly contiguous human genome assemblies have been wildly successful in advancing our understanding of constitutive heterochromatin, finding heterologous recombination between acrocentric chromosomes (Guarracino et al. 2023). Although long-read genome assemblies of the quality of *T2T-CHM13v2.0* assembly are still unfeasible at large population scales, this resource is nonetheless useful for population genomic analyses by improving variant calling and alignment, which we can leverage in the analysis of satellite evolution (Aganezov et al. 2022).

In order to study the variation of simple satellites in human populations and take advantage of the large number of public human NGS libraries, we employ alignment-free methods that circumvent issues of aligning repetitive short-reads and apply them to 2,504 high coverage (∼30x) libraries from the 1,000 Genomes project (Byrska-Bishop et al. 2022). We use *k-Seek*, a method that mines unassembled short-reads for tandemly repeating k-mers 1-20 bp long, to quantify the abundance and variation of simple satellites in these diverse human populations (Wei et al. 2014). We find that the simple satellite content in humans is dominated by repeats of ≤ 5 bp, with the most abundant simple satellites being homopolymers (A/T and C/G) and *Hsat3*. We further use statistical modeling to find interspersed and covarying blocks of AT- and AG-rich repeats, informing us of the mutational processes generating satellite arrays. We also leverage the *T2T-CHM13v2.0* genome assembly and SNPs from the 1,000 Genomes Project to make inferences about the structure and relatedness of centromeric simple satellites. We show that centromeric variation is structured and is predictive of centromeric satellite abundance. Overall, this work integrates short-read data and the *T2T-CHM13v2.0* contiguous assembly in a novel way to describe hitherto unknown variation of human simple satellites.

## Results

We analyzed the tandem repeats in 2,504 human genomes from unassembled short-read libraries sequenced to ∼30x depth from 1,000 Genomes Project using *k-Seek* (Wei et al. 2014; Byrska-Bishop et al. 2022). *k-Seek* mines short-read libraries for repeating monomers of 1-20 nucleotides length that are arrayed in tandem within each read and outputs the identity and number of copies of each repeat found.

### Short-monomer satellites dominate simple satellite content

We discovered 54,353 tandem repeats, with most having low abundance. After correcting copy-number estimates for read depth and GC bias and only including tandems that have at least 1 kb abundance in at least one individual, 126 tandem repeats remained. Additionally, we only consider a tandem repeat to be present in an individual if it spans at least 200 bp. This filtering was done to enrich for long arrays of tandem repeats that are more likely to be derived from heterochromatin rather than microsatellites. Notably, this filtering excluded all 13- and 14-mers from our analysis due to their low abundance (Figure 1a). We also ran *k-Seek* on the *T2T-CHM13v2.0* genome assembly fragmented into 150bp long segments analogous to single-end reads and found only 5% of the total tandem repeats from the 1,000 Genomes Project (2,717 tandem repeats vs. 54,353 tandem repeats) and only 107 of the simple satellites in the assembly. This suggests that there are tens of thousands of low abundance tandem repeats segregating in human populations that are only discoverable through deep population sequencing. Overall, we find that the abundances of simple satellites in the *T2T-CHM13v2.0* assembly are strongly positively correlated with the mean normalized abundances of the 1,000 Genomes Project (Spearman’s Rho = 0.93, p-value < 1×10^-50^), but the *T2T-CHM13v2.0* assembly had greater abundances of almost all simple satellites, except for homopolymers (Figure 1a). Despite this, the *T2T-CHM13v2.0* assembly had a total of ∼20Mb of simple satellites which is lower than the total simple satellite content of the individuals in the 1,000 Genomes Project (Figure 1b). This may represent differences in the genomes of individuals sequenced or the sequencing platforms used as long-read sequencing has known biases in sequencing simple repeats (Flynn et al. 2020).

**Figure 1.**
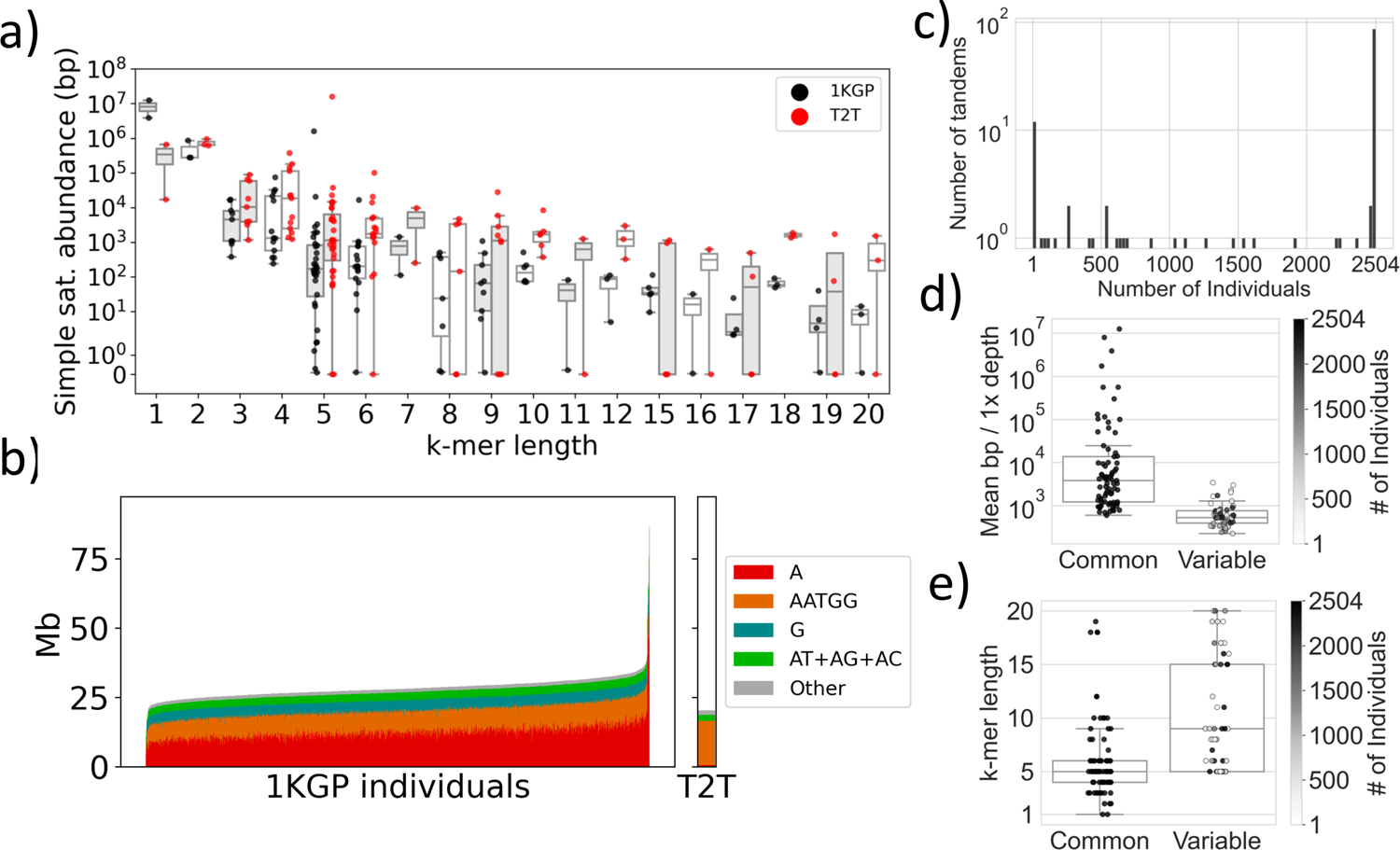
Abundance of simple satellites across the human population. a) Simple satellite abundance for each simple satellite (k-mer) is plotted as the number of basepairs normalized to 1x sequencing depth for the 1,000 Genomes Project (“black”: 1KGP), or the total number of basepairs for each simple satellite in the *T2T-CHM13v2.0* assembly (“red”: T2T). Each point represents a simple satellite, ordered by the monomer length, *k*. Boxes are colored in with white or gray to aid visualization. b) Barplot showing the total simple-satellite content of each of the 2,504 individuals in the 1,000 Genomes Project (1KGP) and the *T2T-CHM13v2.0* assembly (T2T) coloring in the six most abundant simple satellite sequences: A, *Hsat3* (AATGG), G, AT, AG, and AC. The remaining 120 simple satellites are represented by, “Other”. Individual bars are ordered by increasing total repeat abundance in Mb normalized to 1x depth for 1KGP and total Mb for T2T. c) The simple satellite frequency spectrum, which shows presence or absence of a simple satellite in at least 200 bp of abundance across individuals. The vast majority of satellite sequences are present in all 2504 individuals (“Common”), with the left-hand tail representing satellite sequences that are present in a subset of the human population (“Variable”). d) Boxplot showing the mean abundances in basepairs normalized to 1x depth of satellite sequences present in all individuals (“Common”) and those present only in a subset of individuals (“Variable”). Each dot represents a simple satellite sequence and is colored by how many individuals the satellite is present in. “Common” satellites have greater mean abundances than “Variable” satellites (Mann-Whitney U; p-value << 0.05). e) Boxplot showing the monomer length (*k*) of satellite sequences present in all individuals (“Common”) and those present only in a subset of individuals (“Variable”). Each dot represents a simple satellite sequence and is colored by how many individuals the satellite is present in. “Common” satellites have shorter monomer lengths than “Variable” satellites (Mann-Whitney U; p-value << 0.05).

We found that shorter monomers tend to be more abundant than longer ones (Figure 1a). With some exceptions including the pentameric repeat AATGG that is an extreme outlier, there is a decreasing relationship between the average abundance of tandem repeats and their monomer length (Figure 1a). We find that the A/T homopolymer accounts for over half of the simple satellite content in individuals, approximately 12 Mb on average (Figure 1b). The second-most abundant monomer is the pentameric monomer, AATGG (∼1.5 Mb on average), which is the primary repeat unit of the centromeric *Hsat3* satellite (Prosser et al. 1986). Those two repeats plus the G/C homopolymer and the dinucleotide repeats, AT, AG and AC, make up ∼98% (∼19 Mb) of the average repeat content (Figure 1b). The abundance of short satellite repeats may suggest shorter sequences are more easily amplified during replication than longer ones. However, A/T homopolymer abundance also results from retrotransposition of poly-adenylated mRNAs, including from transposable elements, that then expand via replication slippage or unequal exchange (Nadir et al. 1996; Ahmed and Liang 2012).

A/T abundance appeared highly variable across individuals, however this variation is likely largely technical. Ten individuals have extraordinarily high A/T homopolymer content ranging between 30 Mb to 70 Mb, over twice as high as the average (Figure 1b), but eight of these ten individuals had libraries sequenced on the same sequencing run (Supplemental Table 1). After investigating further we find that 2-color sequencer platforms have correlated GC-biases in k-mer content within sequencing runs which can account for large portions of the variation in satellite abundance across libraries. In the case of A/T, the variance explained by the sequencing run is ∼20% and therefore it is likely that these extremely long runs of A/T are a technical artifact (Supplemental Figure 2c). We describe the identification and modeling of these technical biases in full in the *Methods* (*Identification and modeling of technical artifacts of sequencing run)*.To account for these technical effects we directly model sequencer run as a covariate in our statistical models.

We next asked whether there are tandem repeats that are private to specific populations or sets of individuals. To do this we count the presence or absence of each tandem in every individual and use this to compute a satellite-frequency spectrum, a similar idea to the allelic site-frequency spectrum, where we simply report the presence or absence of k-mers across all individuals (Figure 1c). We find that 77 tandem repeats are present in all individuals, which we call “Common”. The remaining 49 tandem repeats are found in a variable number of individuals. These “Variable” tandem repeats range in frequency from singletons and doubletons to being present in all but one individual (Figure 1c). The “Common” tandems tend to be more abundant in individuals and have shorter monomers than “Variable” tandems (Mann-Whitney U; p-value << 0.05) (Figure 1d,e). Despite the lower average abundance of the “Variable” tandems we surprisingly find four singletons that have abundances between ∼5-700 kb in those rare individuals. We reiterate that tandems were removed from individuals if their abundance was below 200 bp; thus some additional individuals may have small amounts of these tandem repeats.

### Mutation-step networks identify cliques of highly similar monomers

To assess the relatedness of satellite sequences we performed pairwise global alignment of all monomers with a length > 3. We then generated a graph such that each node represents a monomer and each edge represents one mutational step. We define a single mutational step as either a single nucleotide difference or a single contiguous indel. This produces a highly connected network with only one node (AAGAAGAAGGAAGAAGCACG) that is disconnected from the larger graph (Supplemental Figure 4). We then employed a Leiden community detection algorithm to find tightly connected “communities” of monomers that are similar in sequence and therefore may have shared ancestry which we dub “cliques” (Traag et al. 2019). Given the short length of these sequences it is not possible to exclude the possibility of similar monomers emerging convergently, nevertheless it is a useful approach for identifying satellites that are candidates for being derived from a common ancestor. Using this approach we identified five cliques of satellite monomers with similar sequence composition.

We identify three cliques of satellite sequences that are generally enriched for specific bases, the AT-rich, AG-rich, and AGAT-rich cliques (Table 1). The two remaining cliques are instead defined by a few abundant monomers. For example we find a clique that includes *Hsat2* (AATCGAATGG) and *Hsat3* (AATGG) that may be derived from the AATGG base monomer, which we dub the *Hsat2/3* clique. We also find a clique containing the highly abundant telomeric repeat (AACCCT) (Table 1).

The *Hsat2/3* clique has the greatest total mean abundance (∼8 Mb) driven by the highly abundant *Hsat3* repeat followed by the AG-rich (∼250 kb) and AT-rich (∼250 kb) cliques.

### AG- and AT-rich repeats form interspersed cliques of co-varying abundance

Previous work studying the population and interspecific variation of simple satellites observed correlated abundances, which may be attributable to concerted amplification of satellite sequences (Wei et al. 2014; Flynn et al. 2021). In order to explore the possibility of concerted amplification of satellites in human populations we modeled the relationship between two satellites under a negative binomial linear model such that,

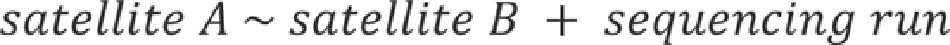

Where, the abundances of satellites “A” and “B” represent the copy numbers of any two satellites and sequencing run is a covariate to model batch effects. From this model we report the proportion of variance in satellite A copy number explained by satellite B (*R^2^*), quantifying the strength of the relationship in copy number. We computed this model for the 77 satellites that were present in all individuals (Supplemental File 3).

We find that although on the whole satellites are not predictive of each other’s abundance (mean *R^2^* = ∼0.07), they are predictive among members of three of the satellite cliques: the AT-rich clique (mean *R^2^* = ∼0.11), the *Hsat2/3* clique (mean *R^2^* = ∼0.16), and the AG-rich clique (mean *R^2^* = ∼0.17) (Figure 2a). To more formally determine which cliques are enriched for covariation in satellite abundance we calculated a Silhouette Score, a score that reflects the quality of the inferred satellite cliques. We first inferred a correlation coefficient by taking the square-root of the *R^2^* value and inferring the sign of the relationship from the beta coefficient. We then computed Silhouette Scores for each clique using the pairwise correlations, which represent the degree of positive relationship of satellites within each clique (Rousseeuw 1987). Silhouette Scores are a useful summary statistic because they are weighted by the overall correlation structure of the data and are bounded between 1 and −1. This means that a positive Silhouette Score represents a clique where satellites have high inter-clique positive correlation and low intra-clique positive correlation, while a negative score represents the opposite. We find three cliques with modestly positive average Silhouette Scores: the AT-rich clique (Silhouette = ∼0.10), the *Hsat2/3* clique (Silhouette = ∼0.07), and the AG-rich clique (Silhouette = ∼0.07). We suggest that members of these cliques may share common origins and are being co-amplified across individuals. In contrast, the telomeric-containing clique and AGAT-rich cliques have negative Silhouette Scores of approximately −0.1 and −0.07, respectively.

**Figure 2.**
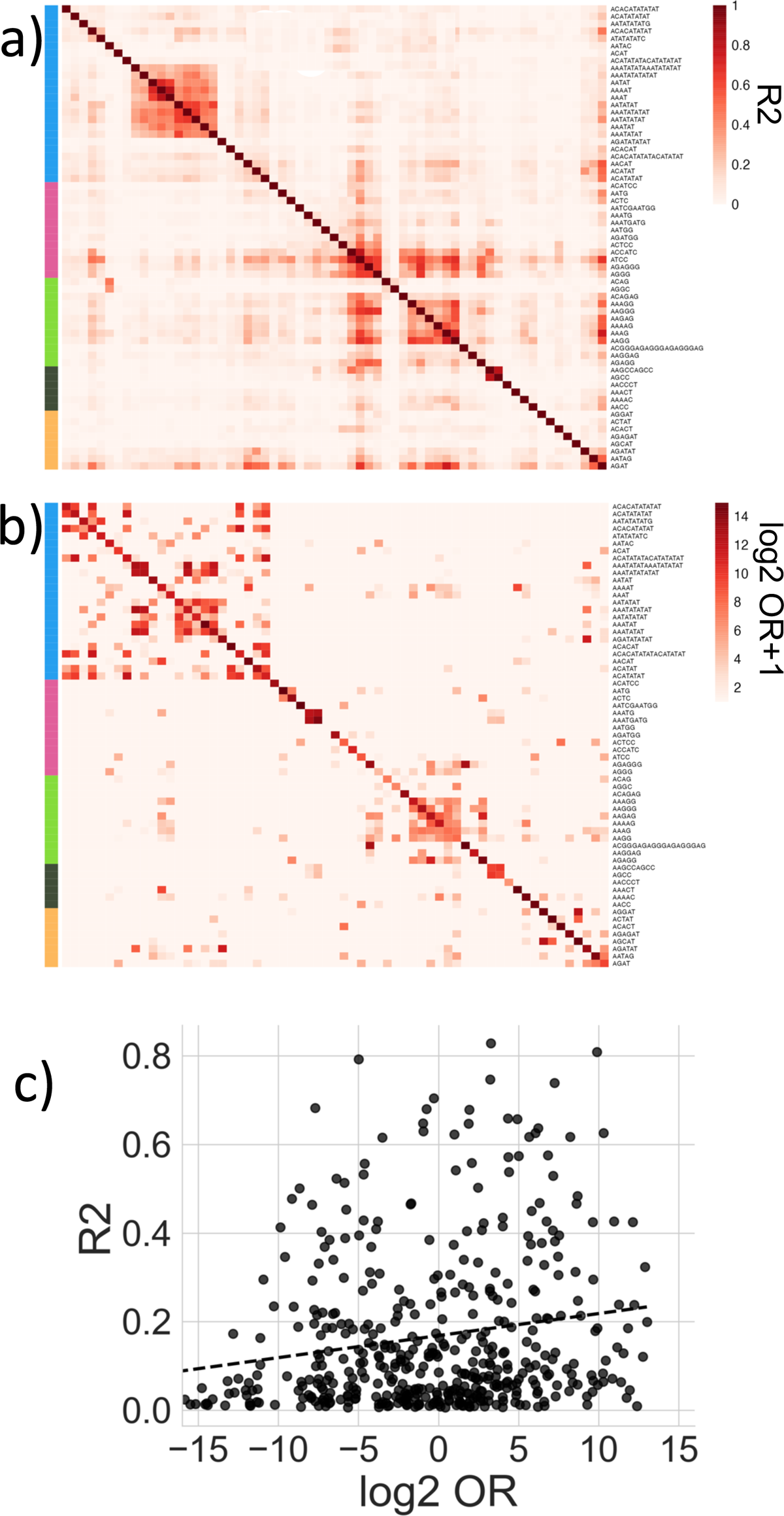
Relationship of covariation in copy number and interspersion of simple satellites forming cliques of similar sequence. a) Heatmap showing the covariation in the copy number between simple satellite sequences as the proportion of the variation explained (*R^2^*). *R^2^* was obtained as the proportion of variation explained by the fixed effects of a negative binomial mixed-effect model. Simple satellites are organized by satellite cliques inferred from the “mutational-step” network analysis: AT-rich (blue), *Hsat2/3* (pink), AG-rich (light green), telomeric (dark green) and AGAT-rich (yellow). b) Heatmap showing the interspersion of simple satellite sequences as the log_2_ of the odds-ratio (OR) + 1. The OR was calculated from the paired-end reads and represents the ratio between the observed number of interspersed reads relative to random chance. Simple satellites as in a). c) Scatterplot showing the positive relationship between the covariation in the copy number (*R^2^*) and the odds-ratio of interspersion (log_2_(OR)) of pairs of simple satellites. Each point represents the pairwise *R^2^* in copy number and the log_2_(OR) for all possible pairwise combinations of simple satellites present in all individuals. There is a positive relationship between *R^2^* and OR (*Spearman’s rho =* ∼0.29; p-value << 0.05). The black dashed line is generated from a linear model for emphasis.

One of the primary ways co-amplification of satellite sequences could occur is through interspersion of satellite monomers. Therefore, we calculated an odds-ratio (OR) of the probability of interspersion of two satellites relative to random chance by taking advantage of the paired-end nature of the data (Figure 2b). Specifically, the OR is the count of read-pairs with two distinct satellites divided by the expected count based on each satellite’s abundance. We compared the pairwise OR of two satellites being interspersed with the *R^2^* of their copy number and found an overall positive relationship with OR and *R^2^* (Figure 2c, *Spearman’s rho* = ∼0.29, p-value << 0.05). Although the correlation coefficient is fairly modest, the pattern suggests interspersion of satellites is contributing to co-amplification. Two cliques have high interspersion OR and *R^2^*. The AT-rich clique has the highest mean interspersion (mean OR = ∼200) and high within-clique *R^2^* (mean *R^2^* = ∼0.11; Figure 2a, 2b). The AG-rich clique is also moderately interspersed (mean OR = ∼66) with high *R^2^* (mean R^2^= ∼0.17). Therefore, these two cliques composed of highly similar AT-rich or AG-rich repeats, respectively, have become interspersed and are forming some higher-order structure, which may lead to co-amplification during replication.

Interspersion is not always related to a covariation in copy number, however. The satellites within the *Hsat2/3* clique covary greatly (mean *R^2^* = ∼0.16), largely attributable to satellites similar to an ACTCC monomer (Figure 2a). However, this clique was not highly interspersed overall, but did have a mean OR of ∼46 due to the extremely high OR between a couple of simple satellites (Figure 2b). Conversely, the AGAT-rich clique has satellites that are highly interspersed (mean OR = ∼213) and yet the average covariation in copy-number within the clique is low (mean *R^2^* = ∼0.06; Silhouette Score = ∼-0.1) (Figure 2a, 2b). These exceptions suggest mechanisms additional to interspersion drive covariation of satellite copy number.

### African population structure of simple satellites driven largely by expansion of AG-rich repeats

Although population structure of human microsatellites has been observed, whether or not this population structure extends to long satellite arrays has not been investigated (Willems et al. 2014). We therefore examined potential population variation and structure in the data by producing a PCA on *z*-score normalized counts of the tandem repeats. We find no evidence of population structure on the first three principal components that account for ∼30.9% of the variation and in fact the first principal component is largely driven by technical variation (Supplemental Figure 1; *Identification and modeling of technical artifacts of sequencing run.)*. However, we find significant clustering of African individuals along PC4 that accounts for 6.8% of the variation (Figure 3a). This African-specific variation is driven by differences in abundance of several k-mers, including ACAG, AAAGGAG, AAAGCAAG, AGATAT and AGAGAT which represent the top five k-mers driving variation on PC4.

**Figure 3.**
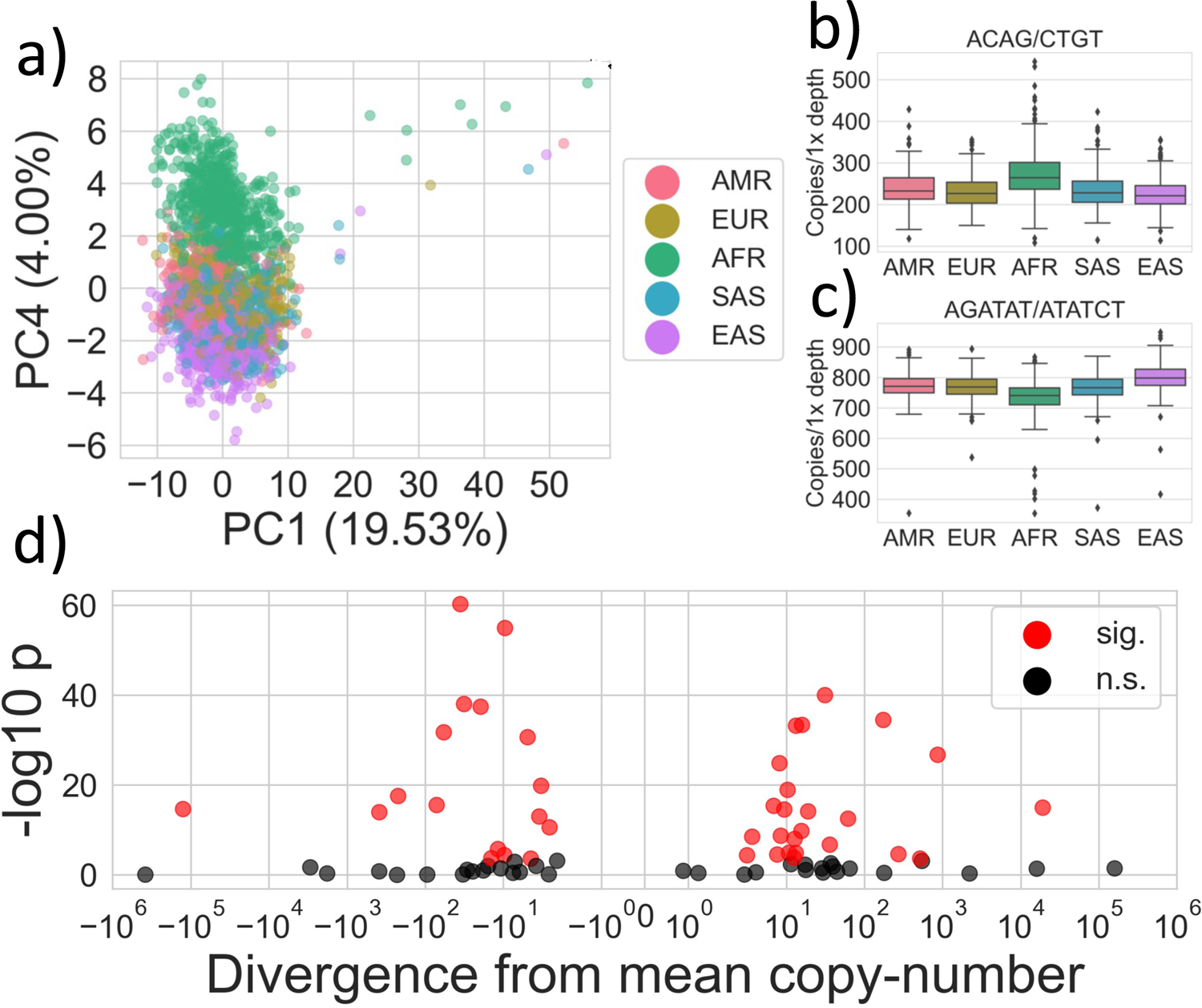
African population structure and differentiation of simple satellites. a) PCA on the normalized simple satellite abundances across individuals. Each point represents an individual in our dataset, colored by their respective “Super Population” (AMR: American, EUR: European, AFR: African, SAS: Southeast Asian, EAS: East Asian). We show PC1, which is driven by technical artifacts (see Results), and PC4, which is driven by African-specific variation in simple satellite abundances. b) Boxplot of the copy number normalized to 1x depth of the ACAG repeat that is higher in African individuals and drives variation on PC4 (same labeling as a)). c) Boxplot of the copy number normalized to 1x depth of the AGATAT repeat that is lower in African individuals and drives variation on PC4 (AMR: American, EUR: European, AFR: African, SAS: Southeast Asian, EAS: East Asian). d) Volcano plot showing the result of a test of population differentiation on simple satellite abundances using a negative binomial mixed-effect model. Each dot represents a simple satellite and reports the p-value and the effect size of the population differentiation as determined by the divergence of the mean copy number of the most differentiated population from the global mean. Each point is colored by whether the p-value was significant given a Bonferroni-corrected critical threshold (⍺=0.05/77).

African populations have on average ∼40-70 additional copies of ACAG, AAAGAG and AAGCAAG and ∼30-60 fewer copies of AGATAT and AGAGAT compared to non-African populations. We show ACAG and AGATAT as representative examples of African-specific population stratification in simple satellites (Figure 3b, 3c). Curiously, the five repeats belong to two cliques, with ACAG, AAAGGAG and AAAGCAAG repeats belonging to the AG-rich clique and AGATAT and AGAGAT to the AGAT-rich clique. The repeats in the AG-rich clique tend to covary in copy number across individuals (mean *R^2^* = ∼0.17) and are interspersed (mean OR = ∼66). Likewise the AGATAT and AGAGAT satellites are also interspersed (OR = ∼20), which suggests that the African population structure is partially driven by divergence in copy number of AG-rich and AGAT-rich higher-order repeats (Figure 3a,b).

We then asked if any other repeats within cliques are differentiated across populations. We implemented a negative binomial mixed-effect model, with population label (AFR, EUR, SAS, EAS, AMR) as a fixed effect and technical artifacts as random effect on 77 of the satellites that are present in all individuals. We find that 44 of these satellites are significantly differentiated and then estimate the effect size of their differentiation by taking the population mean of the most differentiated population and subtracting it from the global mean (Figure 3d). We find that the effect size of the differentiation is quite small in most cases (median effect size = ∼+/-13 copies), however there are a few exceptions. For *Hsat3* (AATGG), East Asian individuals have ∼125,000 fewer copies than the global mean (Figure 3d, Table 2).

We also find that the AG/CT satellite repeat is highly differentiated with an average of ∼19,000 more copies in African individuals than the global mean. It is striking that this highly abundant AG-rich repeat is also more abundant in African individuals than the rest of the population (Figure 3d, Table 2). Within the AG-rich clique we find eight other repeats that are significantly differentiated. Although three of these repeats are differentiated in African individuals, they are lower in abundance in African individuals than the rest of the population, which is opposite to the trend seen for ACAG, AAAGGAG and AAAGCAAG. The rest of the differentiated satellites within the AG-rich clique are elevated in American individuals and East Asians. Of the AGAT-rich repeats, seven differentiated all with modest effect sizes except for AGAT, being ∼500 copies higher in East Asian individuals (Table 2). Across the other cliques and satellites there are several other signs of modest and strong population differentiation (Table 2). These observations of population-specific satellite variation are consistent with observations of *D. melanogaster* and human microsatellites, although the degree of differentiation is not as strong, which may be due to effects of stabilizing selection on satellite copy number (Stephan and Cho 1994; Wei et al. 2014; Willems et al. 2014).

### Identification of novel telomeric variants and centromeric higher-order structures using the Telomere-to-Telomere assembly

To determine the location of these simple satellite sequences we used BLAST with 20bp long concatemers of the monomers on the *T2T-CHM13v2.0* genome assembly and recorded the location of all significant hits (e-value < 0.01, 100% ID) (Supplemental File 1). 117 out of the 126 satellite sequences were successfully localized to the genome assembly, with the remaining satellites representing “Variable” satellites, many of which were only found in 1 or 2 individuals. Using annotations of the centromere, telomere, pericentromeric region, subtelomeric region and Y chromosome we computed an enrichment score that evaluated how often k-mers were found within these regions relative to the rest of the genome and computed a significance test using a McNemar test with a Benjamini-Hochberg adjustment for multiple testing correction (FDR=5%; Figure 4).

**Figure 4.**
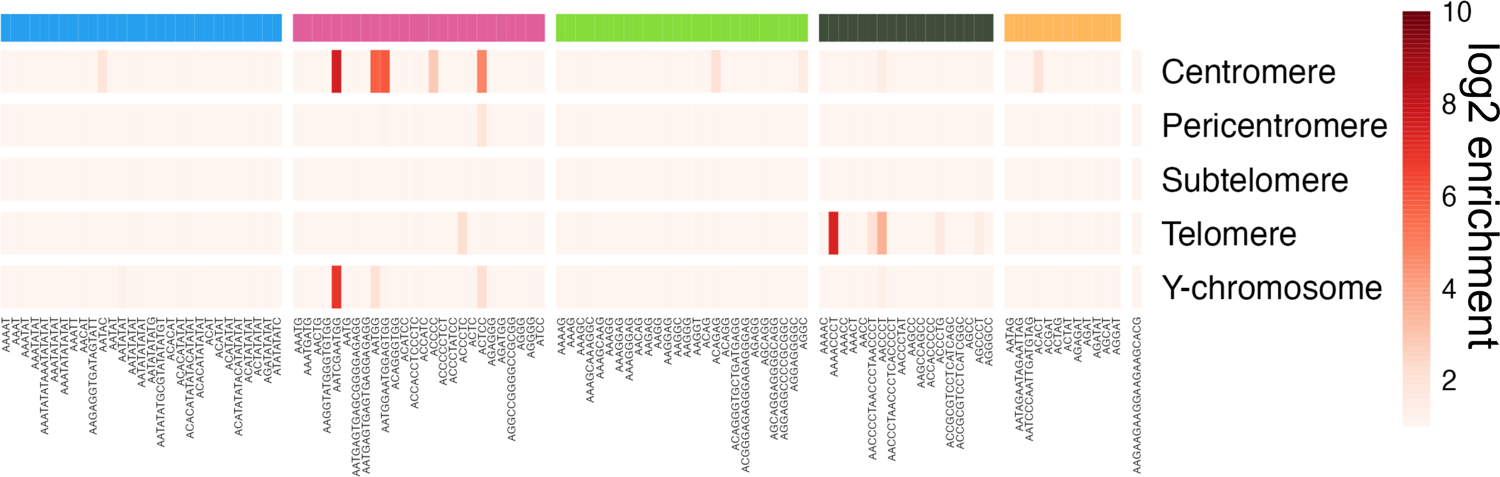
Genomic localization of simple satellite sequences using the *T2T-CHM13v2.0* assembly. Heatmap showing log_2_+1 transformed enrichment scores for presence of a simple satellite in a given genomic compartment. Enrichment scores are the ratio of the number of basepairs of a simple satellite mapping within a given genomic compartment relative to the number of basepairs mapping to the euchromatin as determined by a BLAST search to the *T2T-CHM13v2.0* genome assembly. Simple satellites are organized by satellite cliques inferred from the “mutational-step” network analysis: AT-rich (blue), *Hsat2/3* (pink), AG-rich (light green), telomeric (dark green) and AGAT-rich (yellow).

We found six satellites that are significantly enriched for the telomere. These include the canonical telomeric repeat (TTAGGG) and well known variant telomeric sequences, AGCCCT (CTAGGG) and ACCCTG (TCAGGG) (Allshire et al. 1989; Baird et al. 1995; Alaguponniah et al. 2020). Two others are rare and have not been previously annotated at telomeres.

AACCCCTAACCCTAACCCT (TTAGGGTTAGGGTTAGGGG) is found in 5 individuals and appears to be a higher order repeat structure of the canonical telomeric repeat, TTAGGG, and a novel variant TTAGGGG arrayed to form a TTAGGG(2)TTAGGGG(1) repeating pattern. It is in low abundance in the population (∼500 bp on average in individuals that have the satellite). The AAAACCCT (TTTTAGGG) repeat is found in 530 individuals at extremely low abundance in most individuals (∼200 bp or less), but has a significant expansion in one individual to ∼1 kb. The last telomeric enriched repeat is a known telomeric variant, ACCCTC (TGAGGG) (Allshire et al. 1989; Baird et al. 1995; Alaguponniah et al. 2020). This repeat is clustered with the *Hsat2/3* clique, likely due to its similarity to a common centromeric repeat ACTCC, discussed below.

Within the clique containing known telomere repeats we discovered two other monomers highly similar to telomeric sequences but not found in the genome assembly. We found an exceptionally rare AACCCTAACCCTCACCCCT (TTAGGG(2) GTGAGGG(1)) repeat in two individuals, where the repeat abundance has expanded to ∼200 bp and ∼4 kb, respectively. The AACCCTAT (TTATAGGG) repeat is found in only seven individuals at an average ∼500 bp of abundance and high in one individual of ∼1.3 kb. It is likely that these telomere-like sequences are also found in telomeres, but their rarity in human populations makes finding them in the genome assembly difficult.

We additionally found nine satellite repeats that were enriched for the centromere, but focus discussion on the most highly enriched in the *Hsat2/3* clique. We find centromeric enrichment of the canonical *Hsat2* (AATCGAATGG) and *Hsat3* repeats (AATGG) as well as AATGGAATGGAGTGG, ACTCC and ACCCC. The AATGGAATGGAGTGG repeat is fairly abundant (mean abundance of ∼500 bp) and is present in all but one individual. It appears to be a higher order repeat of *Hsat3* as its repeat structure is AATGG(2)AGTGG(1). Curiously, this repeat is a permutation of CACTC(1)CATTC(2), which includes *Hsat3* (CATTC) and another centromeric repeat, ACTCC (CACTC). The ACTCC repeat has high centromeric enrichment of ∼25x and is fairly abundant, present at ∼13 kb on average. We find that ACTCC has high interspersion (OR = ∼130) with AATGGAATGGAGTGG, and *Hsat2* (OR = ∼37), but not with *Hsat3* (OR < 1) (Supplemental Figure 5). This indicates that *Hsat2*, ACTCC and AATGGAATGGAGTGG form interleaved higher-order structures within the genome. The final highly enriched centromeric repeat is ACCCC which has an enrichment score of ∼6x and is much rarer than the other centromeric satellites, only present in 632 individuals and only at ∼300 bp on average within those individuals. We also find that several of these centromeric enriched repeats are enriched on the Y-chromosome, in particular *Hsat2/3* ATGGAATGGAGTGG and ACTCC. Enrichment of *Hsat2/3* subfamilies on the Y-chromosome was previously observed (Altemose et al. 2014; Altemose 2022).

### Centromeric genetic relatedness explains differences in centromeric satellite abundance

Simple satellites, particularly centromeric simple satellites, are not homogeneously distributed across the genome or centromeres but rather they are enriched within specific centromeric loci (Altemose et al. 2022). We therefore attempted to more finely determine the distribution of the four most highly enriched centromeric simple satellites: AATGG (*Hsat3*), AATCGAATGG (*Hsat2*), ACTCC and AATGGAATGGAGTGG, calculating their relative abundance within specific centromeres using the mapping of satellite monomer concatemers to the *T2T-CHM13v2.0* genome assembly using BLAST. For these centromeric simple satellites we show the proportion of total satellite abundance in the *T2T-CHM13v2.0* genome assembly that is found within each centromere (Figure 5a). We find that ∼52% of *Hsat3* satellites fall within the chromosome 9 centromere, where there is known to be a multi-megabase array of *Hsat3 (Tagarro et al. 1994; Altemose et al. 2022)*. We find most (∼80%) of *Hsat2* satellites within the chromosome 9 centromere and a relatively smaller portion (∼16%) within the chromosome 16 centromere. This is surprising as only chromosome 16 and chromosome 1 centromeres are previously known for having large *Hsat2* arrays (Tagarro et al. 1994; Altemose et al. 2022). The centromeric satellites AATGGAATGGAGTGG and ACTCC have not been previously localized. We find that AATGGAATGGAGTGG is highly abundant in the chromosome 9 centromere (∼66%) and also found moderate amounts in the chromosome 12, 13 and 14 centromeres. ACTCC is almost completely absent from chromosome 9 (1.4%), unlike the rest of these centromeric simple satellites, and instead is fairly evenly distributed across the centromeres of chromosome 13, 14, 15, 21 and 22.

**Figure 5.**
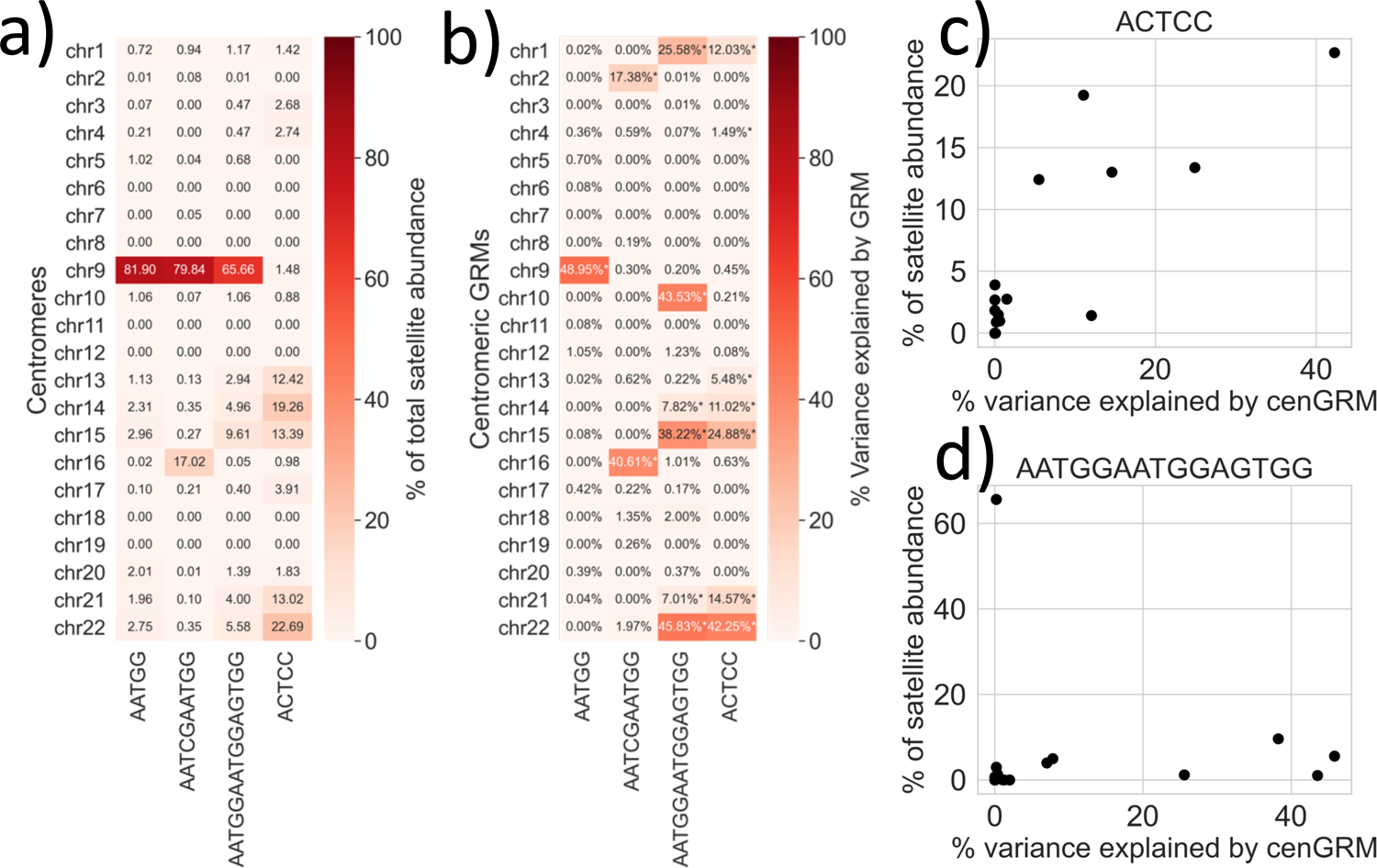
Associations of centromeric genetic relationship matrices. (**cenGRMs) and simple satellite abundances.** a) Abundances of simple satellites that were scored as centromere-enriched within each centromere of the autosomes of the *T2T-CHM13v2.0* assembly as a proportion of the total abundance of each satellite. b) Percent of the variance in centromeric simple satellite copy number explained by the cenGRMs of each autosome. Asterisks represent a significant association that passed a Bonferoni-corrected critical threshold. c) Scatterplot showing the positive relationship of percent variance explained in ACTCC copy number by the cenGRM of each autosome vs. the proportion of ACTCC satellite abundance on each autosome estimated from the *T2T-CHM13v2.0* assembly. d) Scatterplot showing the positive relationship of percent variance explained in AATGGAATGGAGTGG copy number by the cenGRM of each autosome vs. the proportion of AATGGAATGGAGTGG satellite abundance on each autosome estimated from the *T2T-CHM13v2.0* assembly.

One major question regarding satellite evolution is whether the rate at which satellite arrays change in size is much faster than the accumulation of neutral sequence polymorphisms. One way to indirectly answer this question is by asking whether centromeric and pericentromeric genetic variation is associated with differences in satellite copy number. If centromere-linked polymorphisms are predictive of centromeric satellite copy number then this indicates that centromeric satellite copy number evolves at a similar rate to the polymorphisms. We use polymorphisms called from the 1KGP dataset aligned to the *T2T-CHM13v2.0* assembly that are immediately 1Mb flanking and span the centromeres to generate genetic relationship matrices (cenGRMs) (Aganezov et al. 2022). These cenGRMs are representative of centromere-linked genetic variation as the recombination rate surrounding the centromeres is low and linkage disequilibrium is high (Kong et al. 2002; Nambiar and Smith 2016; Langley et al. 2019). Therefore, by using variance-component analysis we can estimate the percent of the variance in centromeric simple satellite copy-number that is explained by the cenGRMs as a way to associate centromeric genetic variation to satellite variation (Figure 5b).

We find that the chromosome 9 cenGRM explains much of the variation of *Hsat3* abundance (∼49%), which is expected given the known presence of the multi-megabase *Hsat3* array spanning the chromosome 9 centromere. The chromosome 2 and chromosome 16 cenGRMs explain the greatest variation in *Hsat2* abundances, of which only the chromosome 16 centromere showed a high proportion of the total *Hsat2* abundance (∼17%). The other two centromeric simple satellites had variances explained that were largely concordant with the centromeric satellite abundances in the genome assembly, with the exception of chromosome 9 (Figure 5b). In fact, we find a strong, positive relationship between the variance explained by the cenGRMs and the proportion of total ACTCC and AATGGAATGGAGTGG satellite abundance in each centromere (ACTCC *Spearman’s rho = ∼0.74, p-value << 0.05;* AATGGAATGGAGTGG *Spearman’s rho = ∼0.61, p-value =0.002;* Figure 5d, 5e).

Chromosome 9 seemed to be a common exception in the association of variance-explained and satellite abundance. Despite finding that 65-80% of *Hsat2* and AATGGAATGGAGTGG content in the *T2T-CHM13v2.0* assembly falls within the chromosome 9 centromere, we find that the chromosome 9 cenGRM explains very little of the variation (∼0.30%) in the copy number of those simple satellites (Figure 5b). We interpret this as indicating the presence of *Hsat2*-like monomers throughout the centromere that were not captured in previous analysis, although it could also reflect errors in the sequence of the chromosome 9 centromere. Other than these exceptions, the greatest variability of these centromeric satellites occurs on the centromeres with largest array sizes. Therefore, centromeric genetic variation is predictive of satellite array size and changes in array size does not outpace the aggregation of genetic polymorphisms.

### Chromosome 9 and 16 cenGRMs show fine-structure associated with Hsat2 and Hsat3

Having noticed that large portions of the variation in centromeric simple satellite abundances can be explained by the cenGRMs we were interested if any of that genetic variation is stratified by population, or other substructure, and if that can explain differences in centromeric simple satellite copy number. We therefore generated a UPGMA dendrogram from all of the cenGRMs to visualize the relatedness of centromeric variation (Supplemental File 4). We find evidence of modest population structure in centromeric variation in several centromeres, particularly of African, East Asian and South Asian variation. However, this is highly dependent on the centromere in question. Chromosome 16 centromeric variation, for example, shows blocks of highly related African and East Asian centromeres (Figure 6a). The chromosome 16 cenGRM is the best predictor of *Hsat2* abundance (*R^2^* = ∼40%), indicating that the population variation of *Hsat2* is likely associated with differences in satellite arrays within the chromosome 16 centromere. *Hsat2* abundance clusters together within some blocks of correlated centromeric variation. In particular, a block of African chromosome 16 centromeric variation (Figure 6a, red block) has high abundance of *Hsat2*.

**Figure 6.**
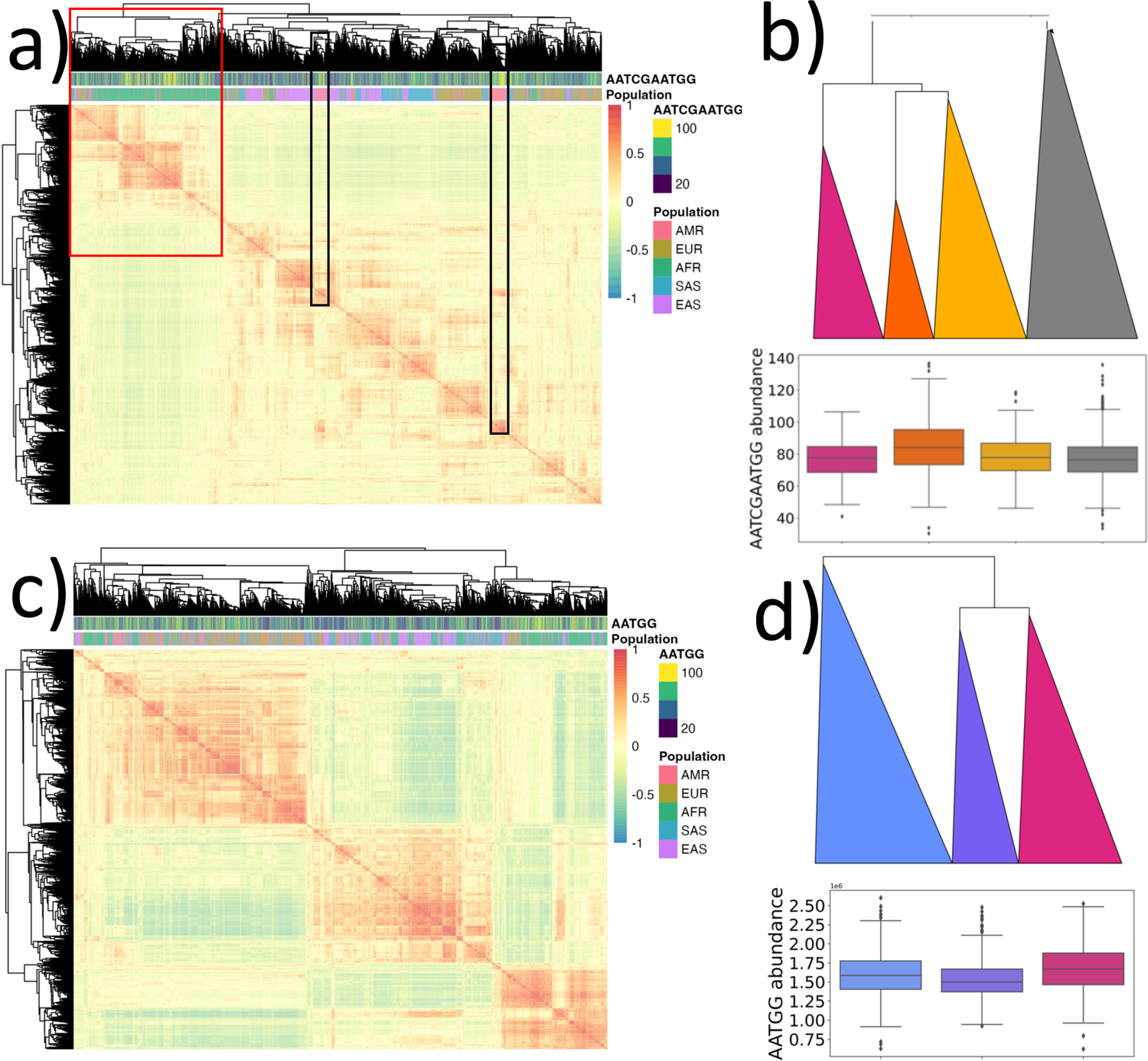
UPGMA clustering of chromosome 16 and chromosome 9 cenGRMs and *Hsat2/3* abundances. a) Clustering of the chr16 cenGRM as a seriated heatmap. Color in cells represents the pairwise correlation in chromosome 16 centromeric genetic variation between individuals. Annotations at the top of heatmap represent the population of each individual (AMR: American, EUR: European, AFR: African, SAS: South East Asian, EAS: East Asian), and the abundance of *Hsat2* (AATCGAATGG) as a percentile. A cluster of largely African individuals is highlighted by the red box and two small clusters of American individuals are highlighted by the black boxes, all of which have relatively high *Hsat2* abundances. b) Collapsed dendrogram of the three highest-level clusters from panel 6a in the red box and the rest of the dendrogram collapsed in gray (top). Boxplot of *Hsat2* (AATCGAATGG) abundances of the three African clusters from the collapsed dendrogram and the rest of the phylogeny in gray (bottom). The orange cluster has significantly higher *Hsat2* abundance (in estimated number of copies) than the rest of the population in gray (Mann-Whitney U; p-value < 1×10^-16^), while magenta and yellow do not (Mann-Whitney U; p-value > 0.05). c) Clustering of the chr9 cenGRM as a seriated heatmap. Color in cells represents the pairwise correlation in chromosome 9 centromeric genetic variation between individuals. Annotations at the top of heatmap represent the population of each individual (AMR: American, EUR: European, AFR: African, SAS: South East Asian, EAS: East Asian), and the abundance of *Hsat3* (AATGG) as a percentile. d) Collapsed dendrogram of the three highest-level clusters from the heatmap of panel 6c (top). Boxplot of *Hsat3* (AATGG) abundances of the three major clusters from the collapsed dendrogram with a y-axis scaled to 1e6. Clusters have variable mean *Hsat3* abundances, with the magenta cluster having the greatest abundance relative to the other two (Mann-Whitney U; p-value < 1×10^-6^).

There is clear evidence of substructure within the primarily African chromosome 16 centromeric variation, with three distinct clusters (Figure 6b). Only one of these clusters of centromeric ancestry is more abundant in *Hsat2* (mean copies per 1x depth = ∼85; Mann-Whitney U; p-value < 1×10^-16^), with the rest having *Hsat2* abundances similar to the rest of the individuals in the tree (mean copies per 1x depth = ∼75) (Figure 6b). When we examine the subpopulation ancestry of the highest *Hsat2* abundance cluster (Figure 6b, orange triangle), we find that individuals with greatest *Hsat2* abundance seem to be of both African ancestry, primarily Yoruban (YRI), Mande (MSL) and Afro-Caribbean (ACB), and American ancestry, primarily Colombian (CLM) and Puerto Rican (PUR) (Supplemental Figure 6). This clustering of American and African chromosome 16 centromeric variation may be indicative of ancestral expansion of *Hsat2* on the chromosome 16 centromere that has become admixed in some American individuals. It should also be noted that there are additional smaller blocks of correlated American centromeric variation with high abundance (Figure 6a black boxes). The subpopulation ancestry of the American individuals in those blocks are predominantly Peruvian (PEL), Mexican (MXL) and Colombian (CLM). We additionally examined the *Hsat2* variation associated with the chromosome 2 cenGRM, but did not find similar patterns, likely because the variance explained by this cenGRM is much lower than chromosome 16 (Figure 5b).

Chromosome 9 centromeric variation is much less structured by population than chromosome 16 and the blocks of correlated genetic variation seem to be much larger (Figure 6c). We observe three major blocks of high correlation that can be divided into three clusters (Figure 6c). These clusters do not have clear population structure, but there are significant differences in *Hsat3* abundance between them (Figure 6d). We find that each cluster is significantly different from each other in *Hsat3* abundance, with the magenta cluster having the greatest abundance of ∼1.67 million copies of *Hsat3* per 1x depth on average (Figure 6d; Mann-Whitney-U p-values < 1×10^-6^). The mean difference between the most *Hsat3* abundant cluster (magenta) and least abundant cluster (purple) is approximately 150,000 copies, which is ∼25,000 copies more than the difference in *Hsat3* abundance between East Asian individuals and the rest of the population. This emphasizes that although chromosome 9 centromeric variation does not have very clear population structure there is still some fine-structure in the centromeric variation that is reflective of the variation of *Hsat3*. And because the chromosome 9 cenGRM was a strong predictor of *Hsat3* abundance (∼50%) and no other centromeres associate with *Hsat3* abundance, it is likely that the most of the variation in *Hsat3* we observe is linked to variation in chromosome 9 centromeres.

## Discussion

### Evidence of high copy number variability of rare, telomeric variants and simple satellites

By examining their presence and absence across human populations we discovered ∼50 simple satellites that are present in only a small number of individuals in the population, including some that have expanded to >1 kb in only one individual. The presence and absence of these satellites appears to not be related to population structure and satellites with longer monomers tend to be rarer in the population. These results emphasize the volatility of rare satellite repeats and mirrors the results of rapid repeat turnover seen in other species (Wei et al. 2018; Cechova et al. 2019 Jul 2). We emphasize an important caveat, that absence of a satellite in an individual does not mean that the monomer sequence does not exist in the genome. Rather it means that the satellite copy number did not pass our detectability threshold of 200 bp. We find that 76 simple satellite monomers, 18 of which are rare in the population, are annotated as microsatellites (Avvaru et al. 2020). Therefore, it is likely that some of these rare satellites are microsatellites that have expanded within a handful of individuals. Large microsatellite expansions doubling the repeat size are not uncommon. (Richard and Pâques 2000). As an example, intergenerational changes in CTG repeats, attributable to myotonic dystrophy, can increase by several hundred copies per generation and reach up to 4,000 copies in length (Tsilfidis et al. 1992; Gomes-Pereira et al. 2007). The expansion may also be driven by replication slippage during cell line maintenance, resulting in a heterogeneous copy number in the cell population that was sequenced. Therefore the copy number estimate is not necessarily representative of every tissue, but the average across the cells sequenced. Additionally, technical artifacts may be biasing our estimates of copy number as well.

We cannot make any inferences as to whether the rare satellite expansions we observe are attributable to any specific microsatellite but using the *T2T-CHM13v2.0* genome assembly we find that among these rare satellite repeats are two telomeric repeats: AAAACCCT (TTTTAGGG) and AACCCCTAACCCTAACCCT (TTAGGG-TTAGGG-TTAGGGG). Variant telomeric sequences have been described previously, particularly TCAGGG, TGAGGG, and TTGGGG. These variant telomere sequences tend to be near the proximal ends of the telomeres and are in high linkage disequilibrium with nearby non-telomeric sequence (Allshire et al. 1989; Baird et al. 1995; Coleman et al. 1999; Baird et al. 2000). The mechanism by which these alternative telomeric sequences are formed and expand is not clear (Allshire et al. 1989). These variants seem to be particularly important for alternative lengthening of telomeres (ALT), a homologous-recombination based mechanism to maintain telomeres that is active in cancers where telomerase is inactive (Henson and Reddel 2010). In cells undergoing ALT, variant telomeric sequences can be found interspersed throughout the telomeres and these repeats, particularly TCAGGG, can recruit nuclear receptor proteins that contribute to increased ALT (Conomos et al. 2012; Alhendi and Royle 2020). The ALT mechanism as well as telomerase-based replication are capable of producing telomeric variant sequences. However, telomerase induced mutagenesis has a preference for generating mutations at the first and third nucleotide while ALT mutagenesis is more stochastic (Lee et al. 2014). The telomeric variants that we have discovered are distinct from previously described variants and are clearly products of indels rather than a nucleotide substitution, which might implicate the more stochastic ALT mechanism. Overall, this discovery emphasizes the high variability of rare satellites, particularly those at the telomeres.

### Population structure of AG-rich and Hsat3 satellites are exceptions to homogenous human simple-satellite content

Population genetic theory predicts homogenization of the satellite arrays within populations due to copy number stabilization of large arrays by selection with some variation driven by copy number mutations and drift (Perelson and Bell 1977; Stephan 1989; Stephan and Cho 1994). Surprisingly, analyses of the tandem repeat content of inbred and experimentally evolved populations has revealed large expansions and contractions of satellite repeats, many of which seemed to be heterogeneously changing across phylogenetic branches (Flynn et al. 2017; Flynn et al. 2021).

However, the dynamics of satellite evolution in natural populations may be quite different due to differences in the selective pressure of inbred and outbred populations. Our results provide the first description of population-specific variation of high abundance simple satellite repeats in human populations, which is quite minor. Only ∼40 simple satellites were differentiated, and the degree of differentiation of most was modest. One major caveat in this analysis is that we are unable to map simple satellite arrays to any particular locus due to the large array sizes. Therefore, satellite abundances are estimated genome-wide from many sources of satellite arrays. If we were able to map the copy number of satellite arrays to discrete loci, we may discover greater differences in the sizes of satellite arrays across populations.

Despite this, these results suggest that in human populations, the most abundant simple satellites retain similar copy numbers, and few satellites have large differences in copy number between populations. The major exceptions are the AG-rich satellites, which are higher in African individuals and *Hsat3*, which is depleted in East Asians. Differences in the evolutionary history of humans may be driving these differences compared to other animals. Human populations have low genetic diversity due to population bottlenecks, which may reflect greater homogenization of satellite arrays (Gravel et al. 2011; 1000 Genomes Project Consortium et al. 2015). Indeed, *D. melanogaster* populations have much greater differentiation in simple satellites than humans, emphasizing the great diversity of simple satellite dynamics across species (Wei et al. 2014).

### Interspersion of satellite cliques of similar sequence partially explains their covariation in copy number

Close sequence similarity of satellites may imply an evolutionary relationship, but because their monomers are such short sequences it is also possible for two similar monomers to appear convergently. A “mutational-step” network-based approach has been successfully used to visualize putative evolutionary relationships of satellite monomers in other species (Flynn et al. 2017; Wei et al. 2018; Flynn et al. 2021). This method does not provide a directed evolutionary trajectory, but rather a structure representing sequence similarity of satellite monomers. Our addition of Leiden clique detection on the “mutational-step” network allows us to identify cliques of highly similar satellite monomers with greater precision (Traag et al. 2019). Analogous clique detection on human pan-genome graphs have provided insight into non-homologous recombination on the short arms of the acrocentric chromosomes (Guarracino et al. 2023).

We successfully identified several cliques of sequences including AT-rich and AG-rich cliques. What is particularly striking about this approach is that by merely placing satellites in cliques by their sequence similarity, we discover that AT-rich and AG-rich satellites covary in copy number across the population and are interspersed. From these observations, we infer that the AT-rich and AG-rich satellite cliques have likely evolved from ancestral AT or AG monomers, respectively, and through repeated expansions have formed complex, interspersed higher-order structures. The interspersed nature of these repeats may be driving their covariation across individuals in the population, whereby unequal exchange amplifies large segments of higher order repeats in tandem (Gripenberg 1964; Craig-Holmes and Shaw 1971; Smith 1976; Perelson and Bell 1977). The relationship between interspersion of similar satellite repeats and their covariation across individuals has been observed in *D. melanogaster* as well, in particular AT- and AG-rich repeats (Wei et al. 2014).

However, much like in previous studies, we find that not all covariation can be explained by interspersion (Wei et al. 2014). In particular, we find that satellites within the *Hsat2/3* clique, specifically monomers that are similar to ACTCC or are AG- and AC-rich, covary in copy number but are not interspersed. The interspersion OR metric is useful at detecting enrichment of many interleaved monomers, but is vastly underpowered to detect a single or small number of junctions between large simple satellite arrays. Therefore, long arrays of monomers that are flanking each other but not interleaved, would show a low OR but could still covary. Such covariation would be stronger if there is linkage disequilibrium between the simple satellite arrays, such as from reduced recombination that occurs at centromeres or on acrocentric arms. It is possible then that discordance between covariation and interspersion is driven by physical linkage of long satellite arrays at low-recombining loci.

### Highly abundant centromeric simple satellites related to Hsat2 and Hsat3 subfamilies discovered mining NGS data

We have identified four simple satellites that are highly enriched for centromeres in the *T2T-CHM13v2.0* genome assembly. Two are the monomers that make up *Human Satellite 2* (*Hsat2*) and *Human Satellite 3* (*Hsat3*) (Prosser et al. 1986; Lee et al. 1997). These centromeric satellites were discovered as satellite bands of variant buoyant density in cesium gradients and eventually had their monomer sequences identified: AATCGAATGG (*Hsat2*) and AATGG (*Hsat3)* (Corneo et al. 1967; Corneo et al. 1968; Corneo et al. 1970; Corneo et al. 1971; Corneo et al. 1972; Prosser et al. 1986). These satellites localize to specific centromeric and pericentromeric regions of the genome, first described using hybridization of fluorescent probes and more recently through bioinformatic analyses of the *T2T-CHM13v2.0* assembly (Tagarro et al. 1994; Altemose et al. 2022). Other work revealed that *Hsat2* and *Hsat3* monomers are also found as components of 24-mer *Hsat2* and *Hsat3* subfamilies, that localize to specific centromeres and pericentromeres (Altemose et al. 2014; Altemose et al. 2022). Our analyses were limited to k-mers up to 20 nucleotides in length, which would exclude those sequences.

However, our approach still quantifies the abundances of the *Hsat2* and *Hsat3* monomers within these 24-mers and by using a naive BLAST approach we manage to localize the monomers to regions of the *T2T-CHM13v2.0* assembly. Our analyses revealed that the *Hsat3* monomer is one of the most abundant simple satellites in the genome and that it localizes primarily to the centromere of chromosome 9. The subfamily *Hsat3B5* is known to dominate the chromosome 9 centromere in a massive 27.6 Mb long array (Altemose et al. 2022; Altemose 2022). The *Hsat3B5* 24-mer is composed of fairly homogenous repeats of the canonical AATGG with a trailing AGCG sequence, which is detected through *k-Seek* and was included in our analysis, although the full 24-mer sequence was not identifiable.

Our approach to localize *Hsat2*, however, produced results that are contradictory to previous literature. We found a great deal of *Hsat2* repeats within the chromosome 9 centromere of the *T2T-CHM13v2.0* assembly, in contrast to previous analyses (Altemose et al. 2022). We are confident that our approach is identifying the correct sequences, because we require an end-to-end alignment of a 20 bp long concatemer of *Hsat2* with 100% sequence identity. The most likely explanation of this discrepancy is that identification of *Hsat2* in the *T2T-CHM13v2.0* assembly previously included sequence information of the 24-mer long subfamilies (Altemose et al. 2022). Therefore, the AATCGAATGG monomer determinant of *Hsat2* may be interspersed within the chromosome 9 centromere, perhaps as a subunit of some other *Hsat3* subfamily, and is only detectable through methods such as ours that directly detect the 10-mer monomer. Additionally, the *T2T-CHM13v2.0* genome assembly may have some degree of sequencing or assembly error, wherein arrays of AATGG are occasionally interpreted as AATCG-AATGG.

*Hsat2* was also found in high abundance within the chromosome 16 centromere, where there is a large *Hsat2B* array (Altemose 2022). Chromosome 2 is also known to have a large *Hsat2B* array, but the *T2T-CHM13v2.0* assembly showed only ∼0.08% of the total amount of AATCGAATGG monomers mapping to the chromosome 2 centromere. We do find that ∼17% of the variation in AATCGAATGG abundance can be explained by the chromosome 2 cenGRM, suggesting that although the abundance of *Hsat2* in this centromere is relatively low, its variability is great (Altemose et al. 2014).

Two other centromeric simple satellites were detected in our analysis, both of which are distinct from previous annotations of *Hsat2/3* subfamilies. AATGGAATGGAGTGG and ACTCC (AGTGG) are highly related to the pentameric monomer that underlies both *Hsat2* and *Hsat3 (Prosser et al. 1986)*. We identify an ACTCC monomer within a few *Hsat3* subfamilies including *Hsat3B1* and *Hsat3B4*, however the monomer does not appear in tandem within these 24-mer sequences, suggesting that we are detecting distinct expansions of the ACTCC monomer (Altemose et al. 2014). Also, detection of this simple satellite in the *T2T-CHM13v2.0* assembly and shows considerable overlap with the known localizations of some of these *Hsat3* subfamilies that contain the ACTCC monomer, particularly on chromosomes 21 and 22 further supporting the hypothesis that the ACTCC monomer has expanded forming its own tandem arrays independent of a subfamily (Altemose et al. 2014; Altemose 2022).

Likewise, the AATGGAATGGAGTGG satellite looks to be a 15-mer subfamily composed of the *Hsat3* monomer and ACTCC, but is not a direct match to any known *Hsat3* subfamily. We do find a highly similar 15-mer monomer within *Hsat3A2*: CATTC-CATTC-GACTC (AATGG-AATGG-AGTCG). This 15-mer could be converted into AATGG-AATGG-AGTGG by simply mutating the penultimate C into a G. *Hsat3A2* also occupies many of the same centromeric localizations as the AATGGAATGGAGTGG satellite, the only exception being chromosome 9 where we find the AATGGAATGGAGTGG monomer in high abundance within the chromosome 9 centromere (Altemose 2022). But we believe that the same sources of error that called into question *Hsat2* abundance in the chromosome 9 centromere are affecting this monomer as well. We contend that our tandem k-mer mining has revealed not only canonical *Hsat2* and *Hsat3* satellite abundances, but also novel centromeric simple satellites that are likely derived or related to existing *Hsat2/3* subfamilies.

### Centromeric ancestry is predictive of centromeric satellite abundance and sheds light on the evolutionary dynamics of centromeres

We find that the genetic relationship matrices generated from centromeric and pericentromeric SNPs (cenGRMs) are predictive of centromeric satellite variation. This indicates that changes in satellite array size are linked to flanking neutral polymorphisms and the rates of change in satellite array size are not substantially different than the gradual accumulation of neutral polymorphisms.

Additionally, the positive relationship between centromeric satellite abundance in the *T2TCHM13.v2.0* assembly and the variance explained by the cenGRM suggests that larger satellite arrays are prone to larger changes in copy number, which would occur if unequal exchange is the primary mechanism that influence array size (Smith 1976). We suggest that centromeric ancestry is strongly predictive of centromeric satellite array sizes. These results are congruent with previous work showing lineage-specific changes in satellite copy number, but here focusing on the centromere scale (Wei et al. 2018; Flynn et al. 2021).

Our analysis of chromosome 9 and chromosome 16 centromeres emphasizes fine-grained structure of centromeric variation and its relationship with *Hsat2* and *Hsat3* abundance. Although in aggregate, *Hsat2* had no evidence of population structure, when visualizing the relationship of the centromeres distinct clusters of largely African and American individuals were found to have greater abundances of *Hsat2*. These individuals seem to share some ancestry at chromosome 16 that is correlated with increased abundance of *Hsat2*. We speculate that African chromosome 16 centromeric haplotypes may be present in the populations of Puerto Rico and Colombia, as these populations are highly admixed with African ancestry (1000 Genomes Project Consortium et al. 2015; Norris et al. 2018). In the case of *Hsat3*, we found that centromeric variation across all individuals could be largely divided into three clusters each with differing mean *Hsat3* abundances. There is no obvious population structure within these clusters, but the mean difference of ∼150,000 copies between clusters is striking and is greater than the differences between populations.

We cannot directly ascribe the changes in *Hsat2* or *Hsat3* abundance to be linked to their respective centromeres. However, by using a linear mixed model we were able to find associations between the cenGRMs and these centromeric satellites, ultimately finding strong associations between genetic variation at the centromere and copy number. From this result we can make the inference that those centromeric satellites are physically linked to centromeres 9 and 16, given that little to no variation was explained by other centromeres. It thus appears that the vast majority of *Hsat2* and *Hsat3* variation is occurring on chromosome 9 and chromosome 16, with the caveat that there may be satellite arrays outside of the centromere that we could not detect. In the cases of the centromeric satellites ACTCC and AATGGAATGGAGTGG, many cenGRMs are strongly explanatory and thus ascribing satellite variation to particular loci is difficult. Although population structure and clusters of genetic correlation were observed for these two centromeric satellites, they are simply too broadly dispersed across the genome for the assumptions of our analysis to hold. Different centromeres bearing these centromeric satellite arrays may be structured in different ways, which would preclude our analysis when aggregating genome-wide satellite abundances.

It should also be noted that these cenGRMs are analogous but not the same as centromeric haplotypes (cenhaps), which have been previously used to infer evolutionary histories, archaic admixture of centromeres, and covariation of *Alpha Satellite* abundances (Langley et al. 2019). Identification of both cenhaps and cenGRMs rely on genetic variation that is linked to the centromere due to low recombination, but our analysis is diploid and unphased (Kong et al. 2002; Nambiar and Smith 2016). Recent study of chromosome X cenhaps in males from the 1,000 Genomes Project discovered centromeric haplotypic variation that is linked to differences in *Alpha Satellite* repeat content and higher-order structures (Altemose et al. 2022). This result correlates well with our finding that a great deal of the variation in centromeric simple satellite abundances can be explained by variation of the cenGRMs. A further important caveat of the cenGRM analysis is that although we can construct UPGMA trees to represent the divergence of the centromeric variation, this is not the same as a tree of phased haplotypes, because the cenGRMs are diploid representations. Despite these limitations we are still able to make inferences on the structure of *Hsat2* and *Hsat3* variation linked to the chromosome 9 and chromosome 16 centromeres without having to map any reads. This form of analysis provides a novel framework for studying repeats and could be extended to the evolutionary study of cenhaps if a reconstruction of the haploid contribution of centromeric satellite content could be performed, such as from single sperm. Additionally, the discovery that linked centromeric genetic variation is predictive of satellite abundance could be extended to predict satellite array size when only SNP data are available.

## Methods

### Tandem repeat identification

2,504 unaligned paired-end fastq files from the 1,000 Genomes Project had their adapters removed and were trimmed of low quality sequence (Q < 20) using *trimgalore*. Trimmed fastqs had repeats identified using *k-Seek*, which identifies tandemly repeating k-mers in each read and quantifies the total abundances of tandem k-mers in each library (Martin 2011; Wei et al. 2014; Byrska-Bishop et al. 2022; Krueger). Optical or PCR duplicate reads containing tandem k-mers were identified by flags from the 1,000 Genomes Project alignment file and removed from the final counts (Byrska-Bishop et al. 2022).

### Tandem repeat identification using k-Seek on the T2T-CHM13v2.0 assembly

The *T2T-CHM13v2.0* assembly was fragmented into 150bp long segments and converted into a fastq format to simulate the generation of single-end reads. However, we did not simulate read depth or any other stochastic sequencing processes. Instead this process generates exactly 1x coverage of the entire genome assembly, precisely capturing the repeat content. We then ran *k-Seek* on these simulated reads to estimate the abundances and identities of tandem repeats in the genome assembly without requiring normalization or correction.

### Estimation of GC-corrected k-mer abundance

We estimated the number of copies of a tandem k-mers in a genome when sequenced to 1x depth by taking the total counts of a particular tandem as calculated by the total number of occurrences in the short-read library normalized by the expected read depth for the tandem’s GC content. To do this we first found all uniquely mappable regions of the *hg38* assembly autosomes using 150 bp long k-mers with 0 mismatches (*k*=150, *e*=0) using *genmap (Pockrandt et al. 2020)*. We next calculated percent GC in 400 bp rolling windows in the uniquely mappable regions using an assembly that was masked by RepeatMasker and excluding any regions with masked positions. 400 bp regions were chosen as they roughly reflect the average fragment length of the libraries (Benjamini and Speed 2012). We next computed the read depth at each position of the hg38 assembly for each library using *mosdepth (Pedersen and Quinlan 2018)*. Using the %GC and read depth information we calculated the expected read depth of positions with a given %GC by binning positions by their corresponding %GC (100 bins representing steps of 1% GC) and taking the average of the read depth of the positions within those bins. We then found what %GC bin each tandem monomer would fall into and divided the tandem’s abundance by the average read depth of that %GC bin, thus obtaining a value that represents the expected abundance, or copy-number, of a tandem when the library is sequenced to 1x depth given its %GC. For %GC bins that had less than 10,000 reads we instead used the median read-depth of bins in the library.

### Statistical modeling of satellite abundance

Satellite abundances were modeled as a generalized negative-binomial mixed-effect model using *glmer.nb(nAGQ=0)* from the R package *lme4 (Bates et al. 2015)*. This model was chosen because it most accurately represents the overdispersed, positive counts nature of satellite abundance. Goodness of fit of the satellite abundance to a negative binomial distribution was done by performing a Kolmogorov-Smirnov test on satellites present in all individuals, which resulted in 46 out of the 77 tested satellites fitting a negative binomial distribution (Supplemental Figure 1a). Of the simple satellites that did not fit the negative binomial distribution only the upper or lower quantiles deviated from the expected values rather than an overall poor fit therefore we concluded it was still appropriate to use a negative binomial model (Supplemental Figure 1b, 1c; Supplemental File 2).

Statistical modeling of this type was only done on satellites present in all individuals in our dataset to simplify the modeling and to avoid using zero-inflated models. Extraction of the variance components of these models was done using the *performance* R package using *r2_nagakawa(by_group=F) (Nakagawa et al. 2017; Lüdecke et al. 2021).* In all cases where statistical modeling of this type was employed, we supplied the sequencer run designation extracted from the headers of the *fastq* files as a random effect to model technical artifacts. Negative binomial mixed-effect models were used to study the covariation of simple satellites, estimation of technical effects and population differentiation.

### Identification and modeling of technical artifacts of sequencing run

We first identified batch effects by visualizing the PCA of normalized satellite abundances (post-GC correction) and coloring by various technical metadata extracted from the fastq headers of our libraries including instrument name, sequencing run and flow cell (Supplemental Figure 2a). We found that several significant outliers on PC1 segregated by these technical data and there was variability along PC1 that was stratified by sequencing run (Supplemental Figure 2b). Additionally, we found that PC1 had a modestly positive, yet significant, correlation with the average autosomal read depth of each library (Spearman’s rho = ∼0.19, p-value < 0.05), suggesting that the read depth may also be playing a significant role in our k-mer abundance estimates. The first principal component is the summary of greatest variability in satellite abundance and explains ∼19.52% of the variation in our data, prompting us to investigate further.

In order to understand how much of an effect read depth and sequencing run have on our estimates of k-mer abundance we employed a negative-binomial mixed-effect model and estimated the proportion of variance explained by library read depth and sequencing run:

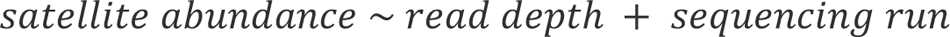

where sequencing run is specified as a random effect in the model, while read depth, computed as the average autosomal read depth, is a fixed effect. For these two models we computed the *R^2^* of the fixed and random effects using Nagakawa and Schiezlith’s method to obtain an estimate of the variation in the abundance of 77 satellites present in all individuals that is explained by technical and biological effects (Nakagawa et al. 2017).

We found that even after correcting for read depth and GC bias there was still significant variation in satellite abundance explained by sequencer run (mean *R^2^* = ∼0.14). The range of *R^2^*was quite wide with ACCATC having an *R^2^* of ∼0.56 while in others, such as ACATATATACATATATAT, the variation explained was close to zero (Supplemental Figure 2c). However, average autosomal read depth did not seem to be a large predictor of satellite abundance (mean *R^2^* = ∼0.005; Supplemental Figure 2c). We find that the satellites that drive variation on PC1 tend to have the greatest variation in their abundance explained by sequencing run, further reinforcing that technical variation is a primary driver of PC1 (Supplemental Figure 2d). Additionally, we find that the *R^2^*of the sequencing run correlates positively with the GC content of the k-mer monomers (Spearman’s rho = 0.36; p-value = 0.001), indicating that the covariation of satellite abundance within the sequencing run may be exacerbated by GC biases (Supplemental Figure 1d). It is possible that we are unable to correct GC bias of high %GC k-mers because there are no uniquely mappable regions of very high GC in the hg38 reference genome to normalize to.

To attempt to correct for the effect of sequencing run on satellite estimates we employed ComBat-seq (Zhang et al. 2020). Although this tool was designed for batch correction of RNA-seq data, the negative binomial linear model employed is appropriate for the positive counts data of simple satellite abundance and the model fit is generally good (Supplemental Figure 1). We employed ComBat-seq with the sequencing run metadata as a batch co-variate and population labels as biological covariates to be retained on the non-GC corrected k-mer counts. We then normalized these corrected counts for read depth and GC bias and employed the same mixed effect model described above to quantify technical artifacts. We found that correcting for batch effects in this way did not significantly change the variance explained by sequencing run (Supplemental Figure 3). We also performed the same analyses on uncorrected and corrected only for read depth and found no significant differences in the *R^2^* (Supplemental Figure 3). Although we are unable to correct these biases directly in the copy number estimates, our identification of the technical artifacts allows us to model these effects directly in linear models adding them in as a random effect.

### Modeling population differentiation and summary statistics

Modeling population differentiation was done using a negative binomial mixed-effect model as described above. The formula used was:

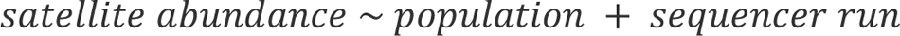

Where satellite abundance represents the abundance in estimated copies normalized to 1x depth, population is a fixed effect representing the population label (AFR:African, SAS:South East Asian, EAS: East Asian, EUR: European, AMR: American), and sequencer run is a random effect representing technical batch. A fixed effect, rather than random effect, was chosen for population labels because of the computational limitations of partitioning variance explained between two different random effects in the negative binomial mixed-effect model (Nakagawa et al. 2017). To our knowledge there are no computationally tractable solutions of partitioning variance explained by two random effects within the same model. Further, we chose levels of the fixed-effect such that they generally reflect the timing of human demographic expansion, *i.e.* African:1, South Asian:2, East Asian:3, European:4, American:5 (Gravel et al. 2011). Therefore, the designations are not completely arbitrary and may have some biological relevance. We use ANOVA on the nested model to determine the effect of population labels on satellite abundance (Bates et al. 2015). To estimate the effect size of the population differentiation of a satellite we simply compute the mean abundance for each population and subtract that from the mean satellite abundance of the entire dataset. We next find which population has the greatest absolute divergence from the mean global abundance and report that value and the population label as the effect size (Table 2).

### Pairwise alignment of monomers and generation of mutational-step network

Monomers were aligned to each other using pairwise global alignment (match = +1, mismatch and gap = 0) taking special care to align both forward and reverse complements of every monomer, as well as all possible offsets (Needleman and Wunsch 1970; Cock et al. 2009). For each pairwise alignment we calculate the number of mutational steps necessary to turn one monomer into another by counting each mismatch or contiguous gap as a single step and call this our “step-score”. This was done to model indels and nucleotide mutations as a single evolutionary step. From all “step-scores” calculated from all possible pairwise alignments of two given monomers we find the minimum “step-score” as the minimum number of mutations to turn one monomer into another. We then generate a network, where each node is a monomer, and draw an edge between two nodes if the minimum “step-score” is 1 (one mutation separates the two monomers), or 0 (they are the same sequence). From this network we implement a Leiden community-detection algorithm to detect “communities” of monomers that are highly connected (similar in sequence), which we refer to as cliques (Traag et al. 2019).

### Calculating interspersion odds ratio from paired-ends

In order to infer whether two, or more simple satellites form interspersed multimeric arrays, we leveraged the paired-end nature of the data and asked how often both reads from a read pair contained a tandem repeat. We queried each read pair flagged by *k-Seek* to contain a tandem k-mer in both mates and added a count to a cell in an *NxN* matrix, where *N* is the number of k-mers in our dataset and each cell represents the number of read pairs with a particular combination of k-mers in both reads. We performed this procedure for each of the 2,504 libraries individually to obtain counts between all k-mers for each individual and also summed across all matrices for an aggregate of all counts for each k-mer combination.

Using these matrices we next asked whether the counts for any given two k-mers would be expected under random chance. To do this we calculated an odds ratio (OR):

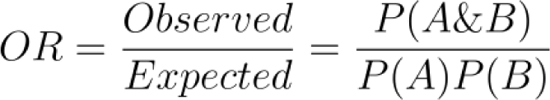

Where we estimate *P(A&B)* as the frequency of paired reads having one mate with k-mer *A* and another mate k-mer *B* and estimate *P(A)* and *P(B)* as the frequency of paired end reads k-mer *A* or k-mer *B* in any mate, respectively. When *P(A&B)* = *P(A)P(B)* then OR = 1, suggesting that the presence of k-mer A and k-mer B are independent of each other. Deviations from this expectation mean that the presence of the two k-mers are not independent, either more likely to be interspersed (OR > 1) or less likely to be interspersed (OR < 1) (Supplemental File 3).

### Localizing tandem arrays to genome assembly using BLAST

To localize satellite monomers to the human genome assembly we used BLAST with the options: *blastn-task ‘blastn-short’ -dust no -soft_masking false*. To localize tandem repeating arrays we generated concatemers of at least 20 bp long and a minimum of two repeats of each satellite monomer sequence and used those as the query sequence against the *T2T-CHM13v2.0* assembly (Altschul et al. 1990; Altschul et al. 1997; Camacho et al. 2009; Nurk et al. 2022). We extracted all BLAST hits that were at least 20 bp in length, had a 100% sequence identity to the query sequence and had an e-value less than 0.01. We then used the coordinates of these BLAST hits and intersected them with the coordinates of the centromeres, telomeres, pericentromeres, and subtelomeres of the autosomes and X-chromosome, as well as the Y-chromosome of the *T2T-CHM13v2.0* genome assembly to determine the localization of these satellites (Supplemental File 2). We defined the pericentromeres and the subtelomere as the region 1 Mb and 10 kb flanking the respective boundary (Supplemental File 2) (Nurk et al. 2022). From those values we calculated a simple score that is the ratio of the total basepairs worth of satellite within the centromere, telomere, pericentromere, subtelomere or Y-chromosome to the number of basepairs that localized to the rest of the genome (Figure 4). We additionally quantified the abundance of satellites within 100 kb bins with 100 kb step-sizes using the same BLAST criterion as above to more finely map the localization of satellite repeats across the genome (Supplemental File 2).

### Generating cenGRMs from 1,000 Genomes Project polymorphism data

In order to generate the centromeric genetic relationship matrix (cenGRM) for each chromosomal centromere we first obtained the Phase 3 VCFs of the 2,504 unrelated individuals in the 1,000 Genomes Project aligned to the *T2T-CHM13v2.0* assembly (Aganezov et al. 2022). This alignment has the benefit of having polymorphisms within the centromeres themselves and certainly flanking the centromeric boundary. We then used these VCFs to generate a GRM using *plink2* including only SNPs and bi-allelic sites with a minor allele frequency greater than 0.05, a minor allele count greater than 80 and a Hardy-Weinberg p-value > 1e-50 that fell within 1 Mb of either side of the centromere boundary: *plink2 --maf 0.05 --hwe 1e-50 --mac 80 --snps-only --max-alleles 2 --make-grm-bin (Chang et al. 2015)*. Centromeric boundaries were defined in the same way as described in the above methods. For chromosome 15 and 16 we used a window of 2.5 Mb flanking the centromere boundary to include sufficient SNPs, such that a statistical model could converge (Supplemental File 4). In general, the boundaries chosen built GRMs from ∼4,000-5,000 SNPs, except for chromosome 15 which included ∼15,000 SNPs.

### UPGMA clustering of cenGRMs

For UPGMA hierarchical clustering we converted cenGRMs, which are covariance matrices, to correlation matrices and used this as a distance matrix for tree construction (Sokal 1958). For determining the distance cut-off for clustering on chromosome 15 and chromosome 16 we manipulated distance cut-offs visually to maximize separation of obvious blocks of highly correlated centromeric genetic variation. This clustering was done primarily to inspect individuals within clusters of interest more closely rather than to make broad claims of clusters of centromeric genetic variation. The cut-off to analyze the chromosome 16 African clusters was 0.12 and to analyze the chromosome 9 clusters was 0.05. As a frame of reference, the mean genome-wide genetic correlation between individuals within the same population is typically between 0.05-0.20, and the mean correlation between two individuals across the whole dataset is approximately 0 (Supplemental File 4).

### Statistical modeling of centromeric satellite abundance and cenGRM variation

We used *GCTA-v.1.94.1* to model the variance in centromeric simple satellite abundance explained by each cenGRM (Supplemental File 4) (Yang et al. 2011). The covariates modeled were: the sex of the individual, first 10 principal components built on all SNPs from the VCF, and sequencing run. These covariates allowed us to control for sex-specific differences, population stratification and sequencing batch effects. GCTA employs a mixed-effect linear regression and estimates the variance explained by the GRM. Although ideally our data best fits a negative binomial model we are limited by the tractability of estimating distinct variance components using a negative binomial mixed-effect model. This model was run on four centromeric simple satellites: AATGG (*Hsat3*), AATCGAATGG (*Hsat2*), AATGGAATGGAGTGG, and ACTCC. We ran a model for each cenGRM for each autosome separately and reported the variance of each centromeric simple satellite that is explained by each autosomal cenGRM and the p-value. Models including all cenGRMs were attempted, but failed to converge in all cases, likely due to a lack of power.

### Data and code availability

For all tandem repeat identification and quantification we used DNA sequences from 2,504 individuals from the Phase 3 of the 1,000 Genomes Project (∼30x coverage Illumina short-read data) (Byrska-Bishop et al. 2022). These short-read data are publicly available within ENA under project accession: PRJEB31736. A VCF file containing alignments of the Phase 3 sequencing of the 1,000 Genomes Project individuals aligned to the *T2T-CHM13v2.0* assembly is available at, https://github.com/schatzlab/t2t-variants (Aganezov et al. 2022). A table containing the sequencing metadata, population of origin and abundances of tandem repeats per individual are available in Supplemental File 1. Results from downstream analyses and associated code are available on GitHub (https://github.com/is-the-biologist/1KGP_SATS).

## Acknowledgements

We thank Elissa Cosgrove and Robert Bukowski for help in the downloading and processing of the 1KGP dataset. We also thank Charles Langley and Sasha Langley for useful discussion of centromeric haplotypes and centromeric satellite variation. This work was supported by the General Medical Sciences Institute of the National Institutes of Health under NIGMS-R01-119125 to D.A.B. and A.G.C.

---

**Table 1.** Satellite abundance of cliques detected using mutational-step network analysis.

Satellites are partitioned into their respective cliques (AT-rich, *Hsat2/3*, AG-rich, Telomeric, and AGAT-rich). One satellite that is disconnected from the network is labeled “NA”. We report the total mean abundance of the clique (“Total”) as well as the mean abundance of each satellite as the basepairs normalized to 1x depth.

**Table 2.** Population differentiation of simple satellite cliques.

We report population differentiation statistics for each simple satellite (“Differentiation”) as the difference in the mean copy number within a population relative to the global mean, and which population is most differentiated (“Differentiated Pop.”). We additionally report the p-value from a negative-binomial mixed-effect model directly testing population differentiation (“p-value”) and whether the p-value is significant under Bonferroni FDR correction (“Significance”; “sig”: significant, “n.s.”: not significant). Satellites belonging to a clique detected from the mutational-step network are labeled as such. Any satellites that are ineligible for any of the categories have an “NA”.

