## Supplemental_Figures for "The structure of simple satellite variation in the human genome and its correlation with centromere ancestry"

### Supplemental Figure 1

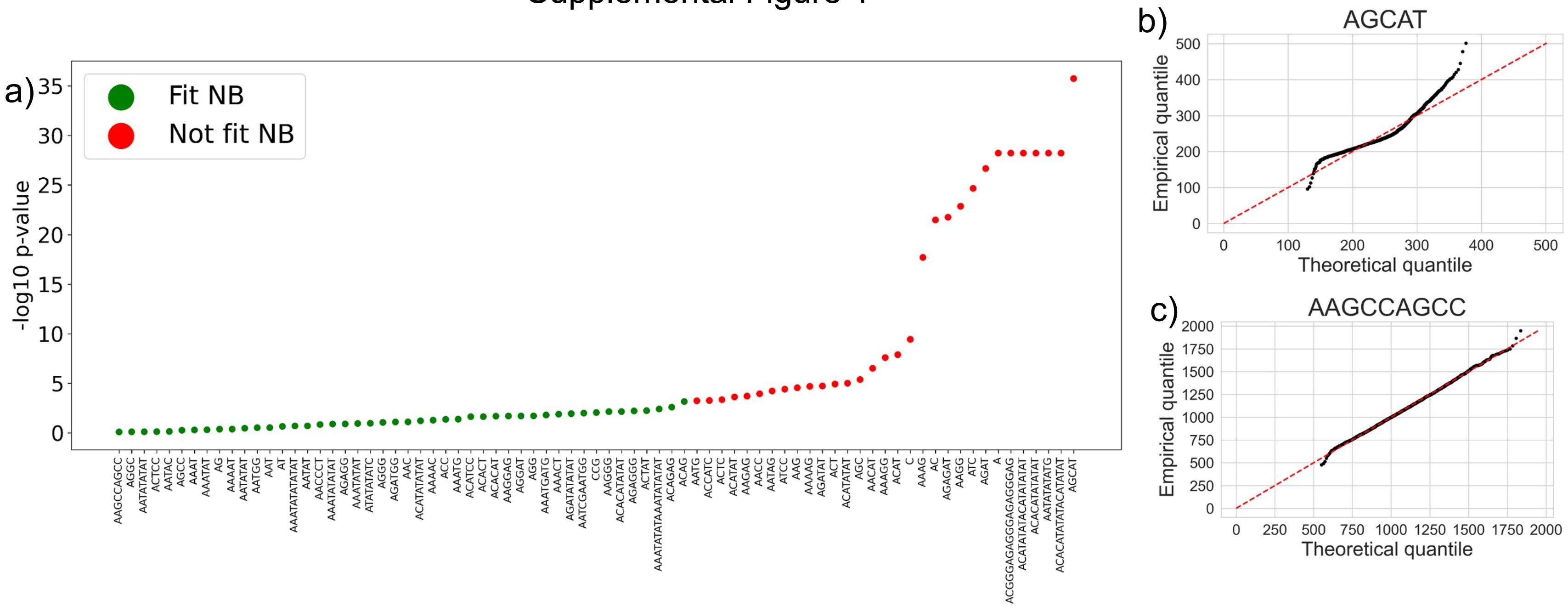

Supplemental Figure 2

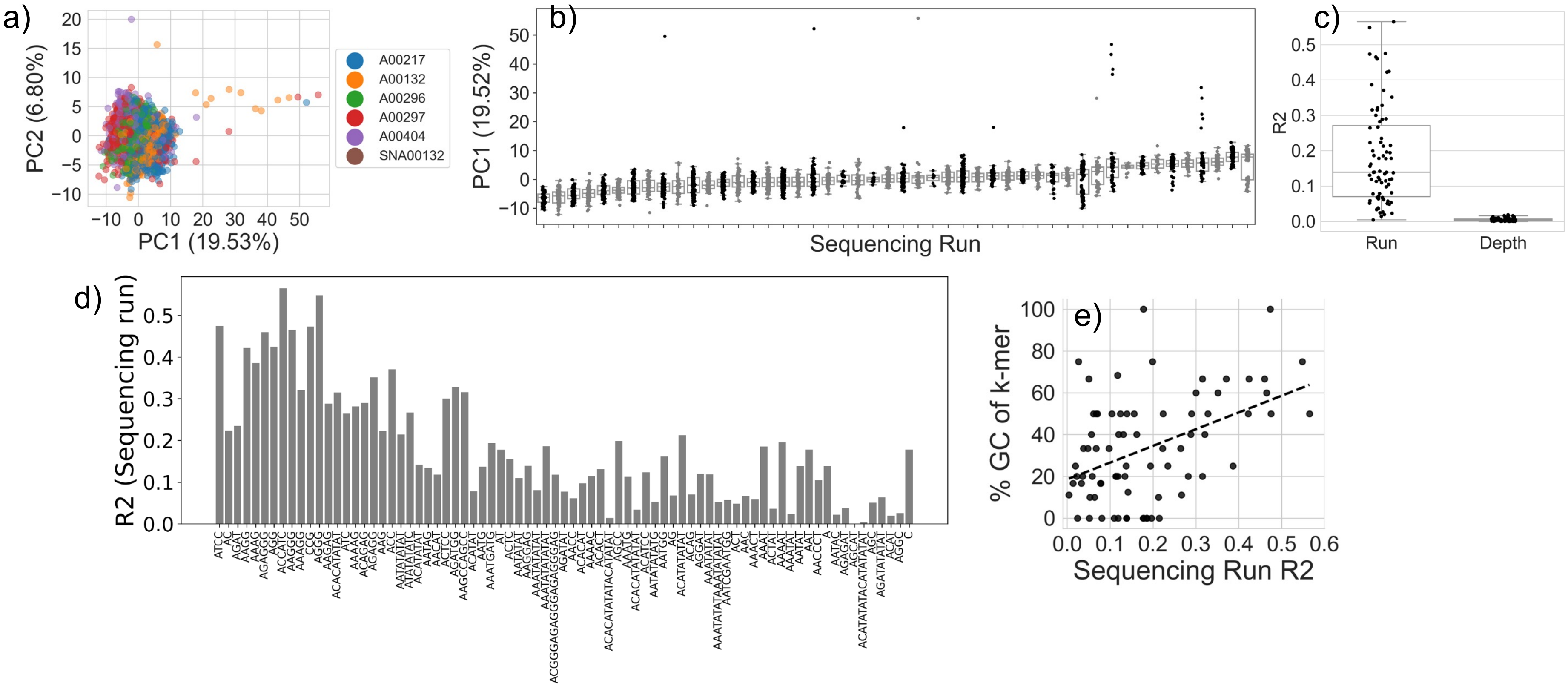

Supplemental Figure 3

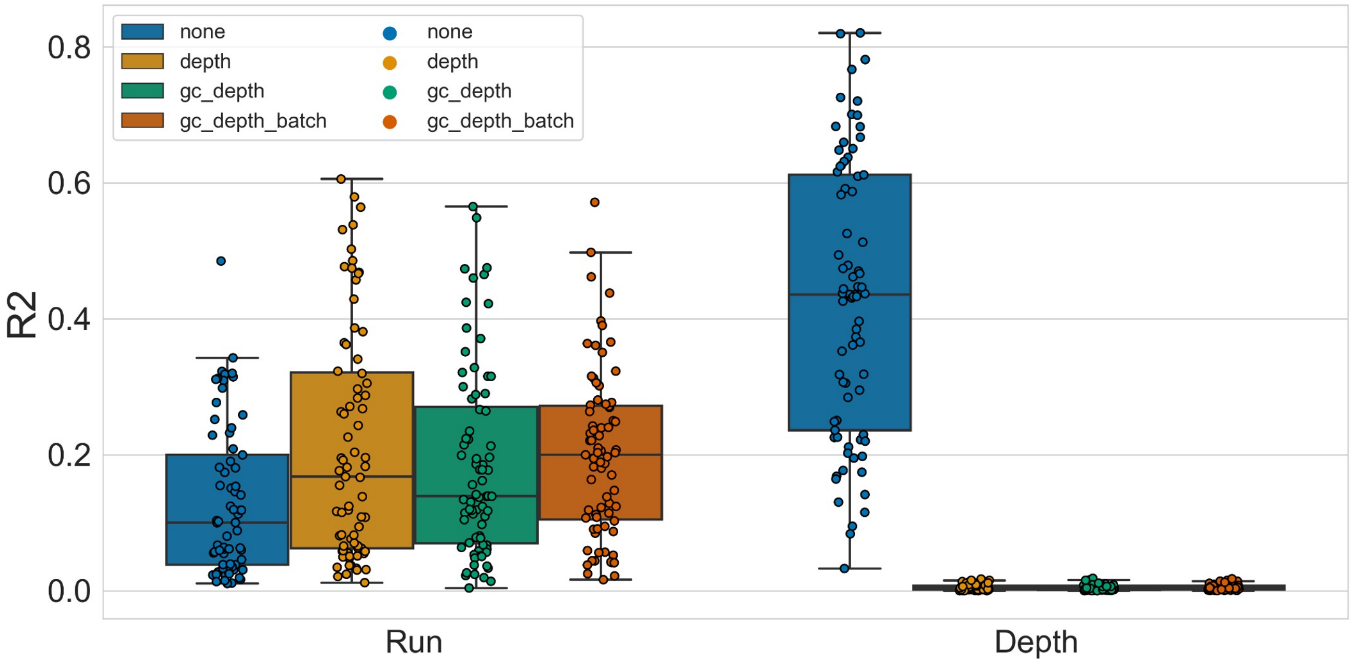

### Supplemental Figure 4

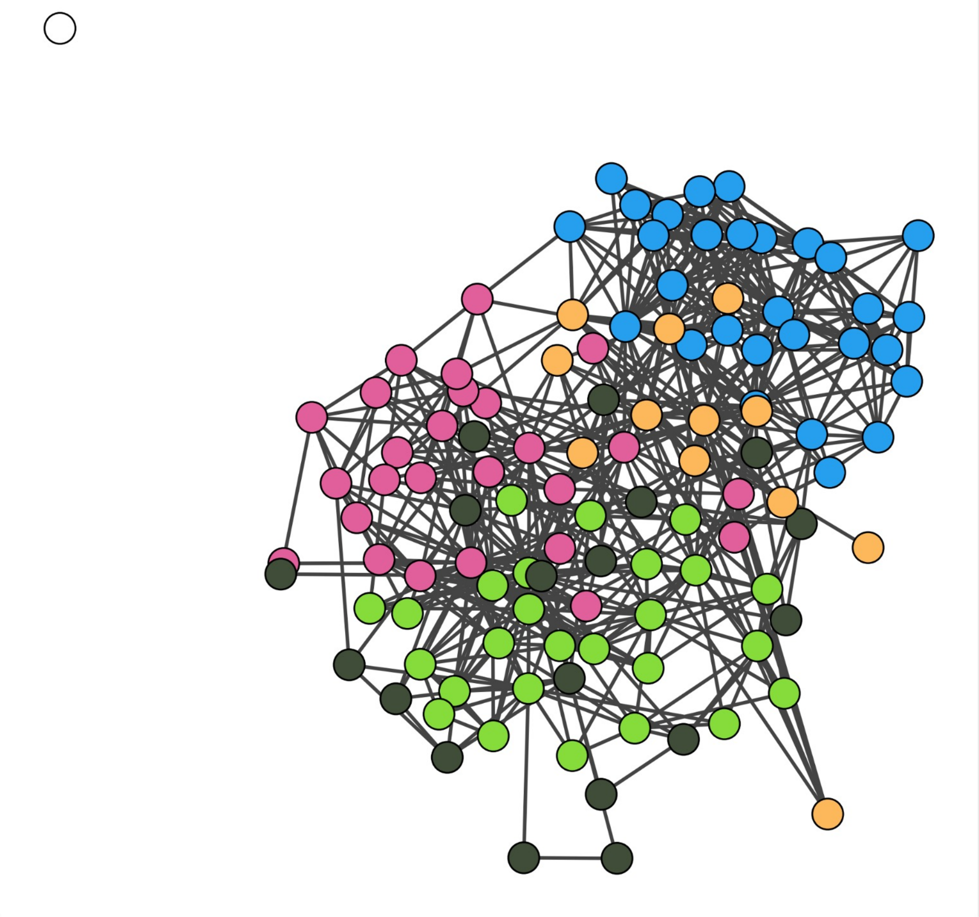

Supplemental Figure 5

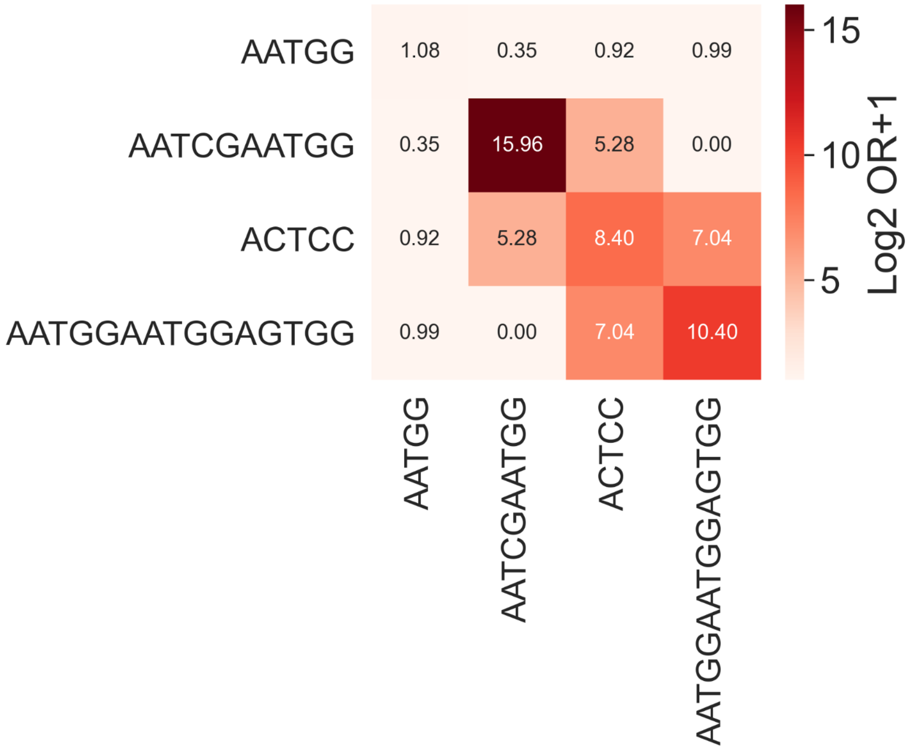

Supplemental Figure 6

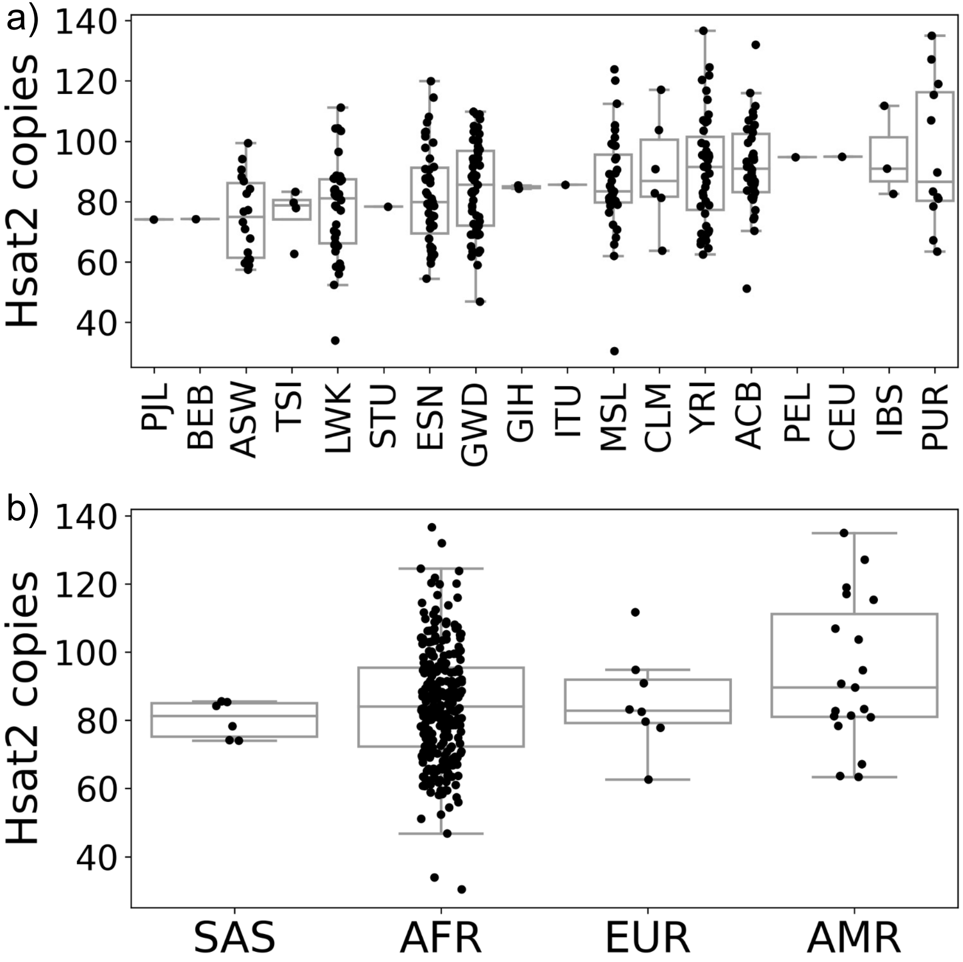

**Supplemental Figure 1. Summary statistics of simple-satellite abundance technical metadata.** a) Result of Kolmogorov-Smirnov test of whether the distributions of simple-satellite abundances fit that of a negative binomial (NB). Each dot represents the result of the test. Low -log10 p-values fit the negative binomial (Green; “Fit NB”), while high values do not fit (Red; “Not fit NB”). b) Quantile-quantile plot showing theoretical and empirical quantiles of the k-mer with the worst fit to a negative binomial distribution panel a). c) Quantile-quantile plot showing theoretical and empirical quantiles of the k-mer with the best fit to negative binomial distribution from panel a).

**Supplemental Figure 2. Analysis of technical batch effects using mixed-effect modeling.** a) PCA of simple satellite abundances showing the first two principal components. Each dot represents an individual and is colored by the sequencing instrument they were sequenced on, emphasizing the substantial effect of technical variation. b) PC1 from panel a) separated by sequencing run. Each dot represents an individual. Data from alternating sequencing runs are colored black and gray to aid visualization. c) The proportion of variance explained (R2) in simple satellite abundance by sequencing run (Run) and by autosomal read depth (Depth) for each satellite. Each dot represents the result of a negative-binomial mixed-effect model used to estimate the contribution of technical effects for each satellite present in all individuals. d) The proportion of variance explained (R2) in simple satellite abundance by sequencing run (Run) ordered by most to least important feature driving PC1. e) Positive relationship between the proportion of variance explained (R2) in simple satellite abundance by sequencing run (Run) and the % GC of the k-mer monomer composing the satellite (Spearman’s rho = 0.36; p-value = 0.001).

**Supplemental Figure 3. Analysis of read, GC and batch correction methods on sequencing run batch effects.** We estimated the proportion of variance explained (R2) in simple-satellite abundance by sequencing run (Run) and by autosomal read depth (Depth) for each satellite under four conditions: No correction (“none”), correction by dividing by autosomal read depth only (“depth”), GC-bias informed read-depth correction (“gc_depth”), and GC-bias informed read-depth correction on ComBat-Seq corrected counts (“gc_depth_batch”). Each dot represents the result of the statistical model for each satellite present in all individuals for the aforementioned data treatments.

**Supplemental Figure 4. "Mutational-step" network of satellite monomers and identified cliques.** Each node represents a satellite monomer and is connected to an adjacent node if one "mutational-step" can transform one sequence into the other. Nodes are colored by their satellite clique identified through Leiden community detection: AT-rich (blue), *Hsat2/3* (pink), AG-rich (light green), telomeric (dark green) and AGAT-rich (yellow). The unconnected white dot, AAGAAGAAGGAAGAAGCACG, does not belong to any clique.

**Supplemental Figure 5. Interspersion of centromeric simple satellites.** Heatmap showing the enrichment of interspersion as an odds-ratio (OR) between centromere enriched simple satellites, AATGG (*Hsat3*), AATCGAATGG (*Hsat2*), ACTCC and AATGGAATGGAGTGG. OR is log2+1 transformed and annotations in cells reflect the transformed value.

**Supplemental Figure 6. *Hsat2* abundances of a differentiated chr16 centromeric cluster by population**. a) *Hsat2* abundance as copies normalized to 1x depth and GC-bias corrected shown for each individual (dot) belonging to the "orange" cluster from Figure 6b separated by subpopulation of origin ("PJL": Punjabi in Lahore, Pakistan, "BEB": Bengali in Bangladesh, "ASW": African Ancestry in SW USA, "TSI": Toscani in Italia, "LWK": Luhya in Webuye, Kenya, "STU": Sri Lankan Tamil in the UK, "ESN":Esan in Nigeria, "GWD": Gambian in Western Division – Mandinka, "GIH":Gujarati Indians in Houston, Texas, USA, "MSL": Mende in Sierra Leone, "CLM":Colombian in Medellín, Colombia, "YRI":Yoruba in Ibadan, Nigeria, "ACB": African Caribbean in Barbados, "PEL": Peruvian in Lima Peru, "IBS": Iberian Populations in Spain, "PUR": Puerto Rican in Puerto Rico). b) *Hsat2* abundance as copies normalized to 1x depth and GC-bias corrected shown for each individual (dot) belonging to the "orange" cluster from Figure 6b separated by superpopulation of origin (AMR: American, EUR: European, AFR: African, SAS: South East Asian, EAS: East Asian).
